# FAF1 and FAF2 enhance human p97-UFD1-NPL4 complex unfoldase activity enabling rational design of p97 activators

**DOI:** 10.64898/2026.02.24.707758

**Authors:** Pritha Dasgupta, Ian R. Kelsall, Gaurav Anand, Anna Pérez-Ràfols, Axel Knebel, Robert Gourlay, Glenn R. Masson, Yogesh Kulathu

**Affiliations:** MRC Protein Phosphorylation and Ubiquitylation Unit, Faculty of Life Sciences, University of Dundee, Dundee DD1 5EH, Scotland, United Kingdom; Division of Cancer Research, School of Medicine, University of Dundee, Dundee DD1 9SY, U.K

## Abstract

VCP/p97 is an AAA+ ATPase that, together with its cofactors UFD1-NPL4 (p97-UN), binds and unfolds ubiquitylated substrates to maintain cellular homeostasis. The human p97-UN complex associates with additional cofactors, but how these cofactors modulate p97-UN activity is not fully understood. Here, we screen for cofactors that enhance p97-UN activity and identify FAF2 as the strongest activator. Using biochemical and structural approaches, we show how FAF2 engages p97-UN and polyubiquitin to promote unfolding. We define a conserved activation motif in FAF2 that contacts both UFD1 and the ubiquitin proximal to the initiator, stabilizing and initiating unfolding in a UFD1-dependent manner. We leverage the features of FAF2 AM to engineer *de novo* proteins that potently enhance unfolding, providing a rational strategy to boost p97 activity. Our findings reveal how cofactors can provide additional adaptive control, fine-tuning human p97 activity to unfold challenging substrates and those modified with short ubiquitin chains.

## Introduction

Efficient proteasomal degradation often requires well-folded proteins to be unfolded and extracted from their native environment. This process is carried out by VCP/p97, an evolutionary conserved AAA+ ATPase, which functions upstream of the proteasome as a segregase and unfoldase (Beskow *et al*, 2009; Olszewski *et al*, 2019). By extracting proteins from membranes, chromatin, and multiprotein complexes, p97 facilitates their subsequent delivery to the proteasome for degradation (Ye *et al*, 2017; Boom & Meyer, 2018). Accordingly, p97-dependent unfolding is essential in several quality control pathways, including endoplasmic reticulum-associated degradation (ERAD), ribosome-associated quality control, and multiple nuclear processes (Maric *et al*, 2014; Brandman *et al*, 2012; Stein *et al*, 2014; Ahlstedt *et al*, 2022). The importance of p97 function is underscored by autosomal dominant mutations that disrupt protein homeostasis and cause multiple neurodegenerative disorders, including multisystem proteinopathy (MSP), amyotrophic lateral sclerosis (ALS), and vacuolar tauopathy (Darwich *et al*, 2020; Watts *et al*, 2004; Johnson *et al*, 2010; Saracino *et al*, 2018).

p97 functions as a homo-hexamer, with each protomer comprising of an N-terminal domain (N) and two tandem ATPase domains (D1 and D2) arranged in stacked rings. The D1 and D2 rings form a central pore through which substrate polypeptides are threaded and mechanically unfolded by ATP hydrolysis-driven conformational changes (Twomey *et al*, 2019; Cooney *et al*, 2023). Substrate recognition is a key regulatory step and is commonly mediated by K48-linked polyubiquitin chains (Richly *et al*, 2005; Blythe *et al*, 2017). This ubiquitin-dependent selectivity is achieved through several adaptor complexes that bind to p97 and recruit specific substrates in different sub-cellular compartments, thereby functionalizing p97 for multiple cellular processes. Of these, UFD1-NPL4 (UN) is a core adaptor complex that contains multiple ubiquitin binding modules, preferentially binding K48-linked chains and recruiting ubiquitylated substrates to p97 (Meyer *et al*, 2000). Both UFD1 and NPL4 bind to the N-terminal domain of p97 via SHP motifs (SHP1 and SHP2) and ubiquitin regulatory X-like (UBXL) domain respectively, with NPL4 arranged in a characteristic tower-like structure above the D1 ATPase ring (Twomey *et al*, 2019; Pan *et al*, 2021a, 2021b). The NPL4 C-terminal domain (CTD) and Mpr1/Pad1 N-terminal (MPN) tower engage distal ubiquitins of substrates modified with K48-linked chains, while the UFD1 UT3 domain binds to proximal ubiquitins located closer to the substrate(Twomey *et al*, 2019; Park *et al*; Sato *et al*, 2019; Williams *et al*, 2023). This multi-site cooperative binding destabilizes one ubiquitin in the chain, the initiator, which becomes captured in the NPL4 groove and unfolds in an ATP-independent manner, positioning its N-terminus for ATP-dependent translocation through the central pore (Bodnar & Rapoport, 2017; Twomey *et al*, 2019; Pan *et al*, 2021a; Ji *et al*, 2022). This substrate engagement mechanism is fundamentally distinct from the proteasome, which initiates unfolding through recognition of unstructured regions in the substrate protein itself (Bard *et al*, 2019).

Beyond the canonical adaptors UN, p97 associates with a diverse array of adaptors which modulate its function and direct it to specific cellular pathways(Buchberger *et al*, 2015; Hänzelmann & Schindelin, 2017). Many of these cofactors are shown to function in concert with UN (Hänzelmann *et al*, 2011; Fujisawa *et al*, 2022). While UN is essential for substrate recruitment and unfolding, the contribution of additional cofactors to p97 function remains incompletely understood. Strikingly, this regulatory cofactor network is far more expanded in human. While yeast Cdc48 functions with roughly 15 cofactors, human p97 interacts with a much broader set of more than 30 cofactors, highlighting increased functional diversification and context-specific regulation. Consistently, comparative studies between yeast Cdc48-Ufd1-Npl4 (Cdc48-UN) and mammalian p97-UFD1-NPL4 (p97-UN) revealed intriguing functional differences. In contrast to yeast Cdc48, mammalian p97-UN requires substantially longer or more complex heterotypic ubiquitin chains to achieve efficient substrate unfolding in reconstituted systems (Blythe *et al*, 2017; Fujisawa *et al*, 2022). Both yeast and human complexes share highly conserved architecture and ubiquitin-binding mechanism (Twomey *et al*, 2019; Bodnar & Rapoport, 2017; Pan *et al*, 2021a), suggesting that any differences in function are unlikely to arise from the core machinery alone. This raises the possibility that mammalian p97 relies on additional regulatory factors, although the identity and mechanisms of such factors remain underdefined.

To address how cofactors impact human p97 unfolding activity, we employ in vitro reconstitution and a focused biochemical screen to identify human p97 cofactors that enhance unfoldase activity of human p97 in the presence of UN. Of the adaptors tested, Fas-associated factor 2 (FAF2/UBXD8), a UBX domain adaptor protein, shows the strongest effect on unfolding by p97-UN. Through biochemical and structural analyses, we define the molecular features that underpin FAF2-mediated stimulation. Based on these insights, we employ computational protein design to engineer *de novo* mini-proteins that mimic FAF2 function within novel protein scaffolds to potently activate p97. Our work reveals how the unfoldase activity of p97 can be enhanced by additional cofactors, an adaptive feature we believe is important to unfold misfolded proteins, and extract proteins from membrane and tightly bound multiprotein complexes.

## Results

### Human p97-UFD1-NPL4 exhibits reduced unfoldase activity

Yeast p97 (Cdc-48) has been well characterized and widely used for biochemical and structural studies(Twomey *et al*, 2019; Bodnar & Rapoport, 2017). In contrast, the regulation and activity of human p97 are not fully understood. To directly compare yeast and human systems, we performed *in vitro* unfolding assays with the yeast and human unfoldase complex using the model substrate monomeric Eos fluorescent protein version 3.2 with N-ter fused ubiquitin (Ub–mEos3.2), hereafter referred to as Ub–Eos (Blythe *et al*, 2017). This substrate mimics ubiquitin fusion degradation (UFD) pathway substrates such as Ub(G76V)–GFP, which has been established as a bona fide p97 substrate in cells (Beskow *et al*, 2009; Johnson *et al*, 1995; Wójcik *et al*, 2006; Dantuma *et al*, 2000; Chou *et al*, 2011). Moreover, Eos undergoes backbone cleavage upon UV irradiation, with a consequent shift from green to red fluorescence. Following unfolding, the two Eos fragments cannot refold leading to an irreversible loss in fluorescent signal. We modified the Ub–Eos with single K48-linked polyubiquitin chains of various lengths: long (Ub^L^, ≥10 ubiquitin), short (Ub^S^, 4-10 ubiquitin), and very short (Ub^VS^, ≤ 4 ubiquitin) (**Figure 1A, EV1B, EV1C**). Following photoconversion, the ubiquitylated Ub–Eos substrates were incubated with reconstituted yeast Cdc48-UN complexes in presence of ATP, resulting in a steady-state decrease in fluorescence that directly reports on substrate unfolding kinetics under single turnover conditions (**Figure 1B**). Human p97 is known to contain multiple post-translational modification sites that are known to influence its adaptor interactions, localization, and activity(Zhao *et al*, 2007; Hänzelmann & Schindelin, 2017). To retain these modifications, the human complex was reconstituted with p97 hexamers purified from a baculovirus expression system (**Figure EV1A**). With insect cell purified human p97 or p97 purified directly from human cell lines (data not shown), we observed a pronounced inefficiency in unfolding activity of human p97 in comparison to the yeast Cdc48 complex. Ub^VS^, substrate modified with less than 4 Ub molecules, was not unfolded by either yeast or human p97, implying that this chain length is below the functional threshold. While the yeast complex efficiently unfolded both short- and long-chain substrates (Ub^S^ and Ub^L^), with unfolding rates of *k* = 0.003 s⁻¹ and *k* = 0.005 s⁻¹, respectively, human p97 displayed markedly reduced activity. Unfolding of Ub^L^ by human p97 (*k* = 0.0008 s⁻¹) was ∼6-fold slower than that of the yeast complex, whereas Ub^S^ was largely resistant to unfolding (**Figure 1C, EV1D**). A similar kinetic difference was observed using Ub_6_–Eos, a substrate bearing a defined K48-linked hexamer, further highlighting the slower activity of human p97 (**Figure EV1E**). This disparity raised the possibility that human p97 may rely on factors beyond UN to achieve efficient substrate unfolding.

**Figure 1:**
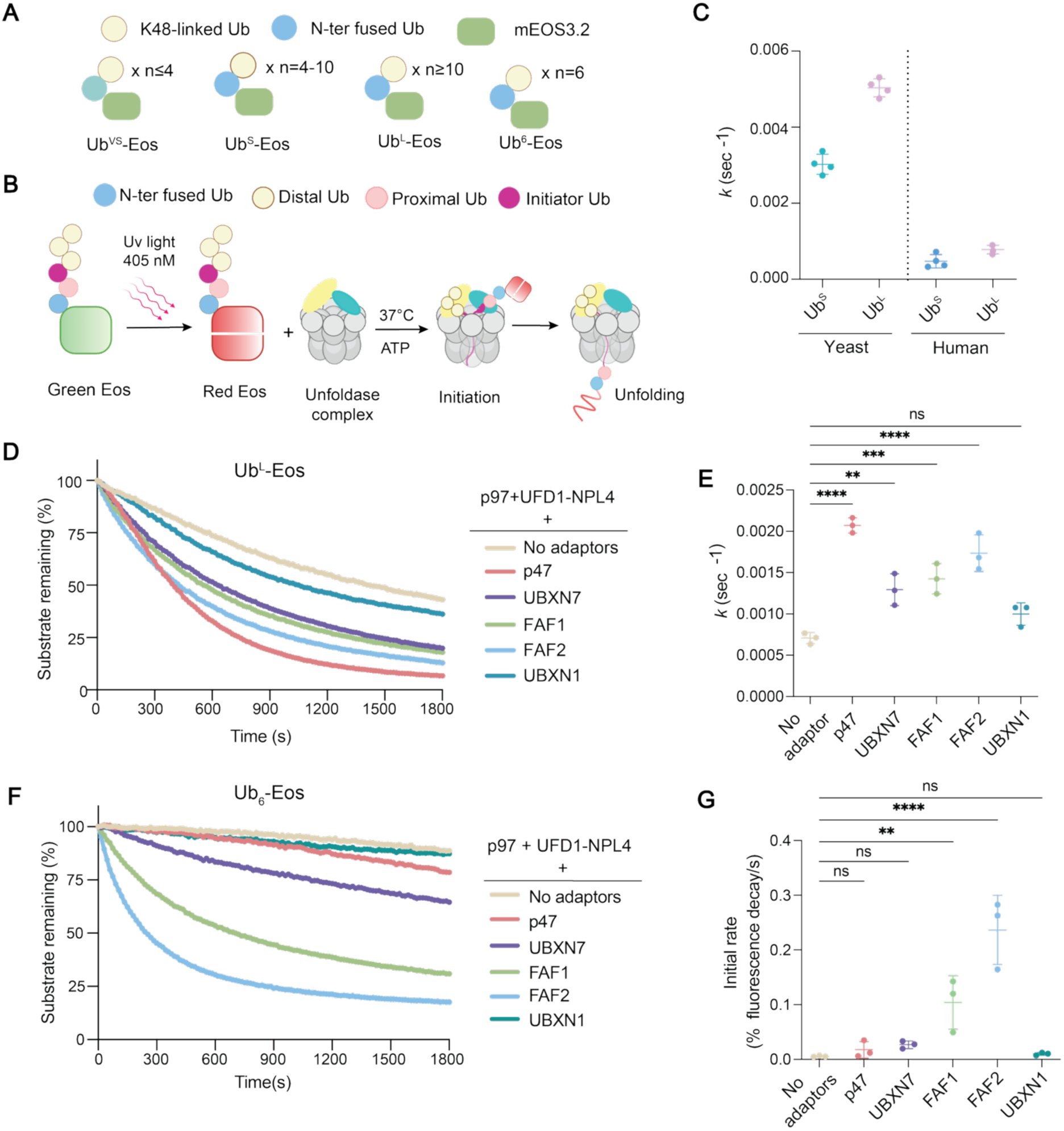
UBA-UBX cofactors enhances unfolding of ubiquitylated substrates by human p97-UFD1-NPL4 complex. (A) Schematic describing the architecture of the ubiquitylated substrate used for the in vitro unfolding assay. Substrate modified with K48-linked ubiquitin chains of ≤4 ubiquitins (Ub^VS^), 4-10 ubiquitins (Ub^S^), and very long chain of ≥10 ubiquitins (Ub^L^). Ub_6_-Eos denotes Ub-Eos substrate modified with a hexameric Ub chain. (B) Schematic representing the principles of substrate unfolding assay with reconstituted p97-cofactor assemblies. (C) Comparison of rate of unfolding (*k*) of Ub^S^ and Ub^L^ substrate by yeast Cdc48– Ufd1–Npl4 (left) and human p97–UFD1–NPL4 (right). Data represent the mean ± SD for n ≥ 3 technical replicates. (D) Single turnover unfolding assay comparing unfolding of Ub^L^-Eos by p97-UN complex alone or upon addition of indicated UBA-UBX adaptors. Data representative of n = 3 independent experiments, each performed with three technical replicates. (E) Comparison of rate of unfolding *k (*sec^-1^) for various p97-cofactor assemblies towards Ub^L^ substrate. Data represents mean ± SD for n=3 independent experiments, each with 3 technical replicates. (F) Screening of the same UBA-UBX cofactors towards Ub_6_-Eos substrate. The unfolding traces are representative of n=3 independent experiments, each with 3 technical replicates. (G) Comparison of the initial rate (first 2 min) of unfolding for various p97-cofactor assemblies towards Ub_6_-Eos substrate. Data represents mean ± SD for n = 3 independent experiments, each with 3 technical replicates. The fluorescence decay is represented by normalizing the fluorescence intensity of each time point to time 0 and to the substrate-only controls. In panel C and E, the unfolding rate *k* (sec^-1^) was determined by fitting the unfolding trace to single exponential decay. In panel G, the initial rate of unfolding was calculated from the linear fitting of the first 2 min of the unfolding reaction because of poor fitting of few conditions into exponential decay function. Statistical significance was calculated by one-way ANOVA followed by multiple-comparison testing (ns, not significant P > 0.05; *P ≤ 0.05; **P ≤ 0.01, ***P ≤ 0.001, ****P ≤ 0.0001).

### Additional cofactors differentially modulate the unfolding activity of human p97

It has been reported that mammalian p97 requires additional adaptors to disassemble CMG helicase modified with short chains (5-7 ubiquitin), whereas such adaptors are dispensable for CMG helicase bearing longer ubiquitin chains (Fujisawa *et al*, 2022). Moreover, proteomics analyses and in vitro studies revealed multiple UBX domain containing p97 adaptors to interact with the UN complex (Raman *et al*, 2015; Hänzelmann *et al*, 2011). Based on these observations, we tested whether UBX proteins that can simultaneously engage ubiquitylated clients influence substrate unfolding by the p97-UN complex. We focused on five p97 adaptors – p47/NSFL1C, UBXN7/UBXD7, FAF1/UBXD12, FAF2/UBXD8 and UBXN1/SAKS1, which all contain a UBX domain and ubiquitin-associated (UBA) domain for binding p97 and ubiquitin respectively. We assessed the ability of these adaptors to stimulate Ub^L^ substrate unfolding in vitro. All adaptors were expressed as full-length proteins, except FAF2, where we deleted the membrane-tethering hairpin (HP) domain to facilitate soluble expression in *E. coli*. We observed a significant enhancement of unfolding activity in the presence of p47, UBXN7, FAF1 and FAF2, whereas UBXN1 showed no detectable effect (**Figure 1D**). Kinetic analysis revealed p47 as the most potent activator among the adaptors tested (**Figure 1E**). Importantly, UN was indispensable, as none of these adaptors by themselves can accelerate the unfolding reaction independently (**Figure EV1F**). These results indicate that p97 together with UN forms the core unfoldase module, while the additional adaptor proteins function cooperatively to modulate or amplify its activity, thereby regulating substrate unfolding.

Model p97 substrates used for in vitro studies are typically modified with very long ubiquitin chains, and it is unclear whether such long ubiquitin chains are made in cells and how prevalent they are. Quantitative proteomics revealed the composition of K48-linked polyUb chain in yeast consists of 2 to 7 ubiquitin moieties per chain at steady state (Tsuchiya *et al*, 2018). Hence, to mimic the predominant pool of p97 clients available in vivo, we generated Ub-Eos modified with K48-linked Ub_6_ (Ub_6_-Eos; **Figure EV1C**), to allow us to assess how a physiologically relevant Ub signal is decoded by the unfoldase complex. The canonical UN adaptor was unable to unfold the short-chain modified substrate, in agreement with the higher ubiquitin threshold of the human p97-UN complex (Fujisawa *et al*, 2022). We next analyzed the impact of UBA-UBX adaptors on unfolding Ub_6_-Eos substrate under identical conditions. While p47 potentiated unfolding of the Ub^L^ substrate, it was unable to support efficient unfolding of Ub_6_-Eos. Of the adaptors screened, only FAF1 and FAF2 supported efficient unfolding (**Figure 1G**). FAF2 showed the most rapid unfolding. In comparison, p47 showed minimal activity and UBXN7 supported only moderate unfolding (**Figure 1F, 1G, EV1G**). In summary, this reveals that adaptor-mediated activation of p97 function varies widely and is strongly determined by the length of the polyUb chain on client proteins.

### FAF2 enhances unfolding of ubiquitylated substrates

Having identified FAF1 and FAF2 as the adaptors that most efficiently potentiate unfolding of substrates modified with both short and long ubiquitin chains, we focused on these two proteins to define the most active p97–adaptor assemblies. FAF2 is a membrane-associated p97 adaptor that plays critical roles in ER-associated degradation, lipid droplet turnover, mitochondrial and peroxisomal quality control pathways (Ganji *et al*, 2023; Koyano *et al*, 2024; Olzmann *et al*, 2013; Zheng *et al*, 2022). AlphaFold3 (AF3) predictions of FAF2 reveal it to contain an N-terminal UBA domain (residues ∼1–55), a short hydrophobic hairpin (HP) region (∼90–122) that anchors FAF2 to the membrane, a conserved thioredoxin-like UAS domain (∼132– 275), a long helical central region (CR) (∼275–347), and a C-terminal ubiquitin-regulatory X (UBX) domain (∼362–445) (**Figure 2B**). DALI searches using either the UAS domain together with the helical region or the helical region alone did not yield any meaningful structural homologues. FAF1 shares a similar overall domain architecture but instead of a membrane-tethering domain, harbours two additional ubiquitin-like domains (UBL1 and UBL2) at the corresponding position **(Figure 2A**).

**Figure 2:**
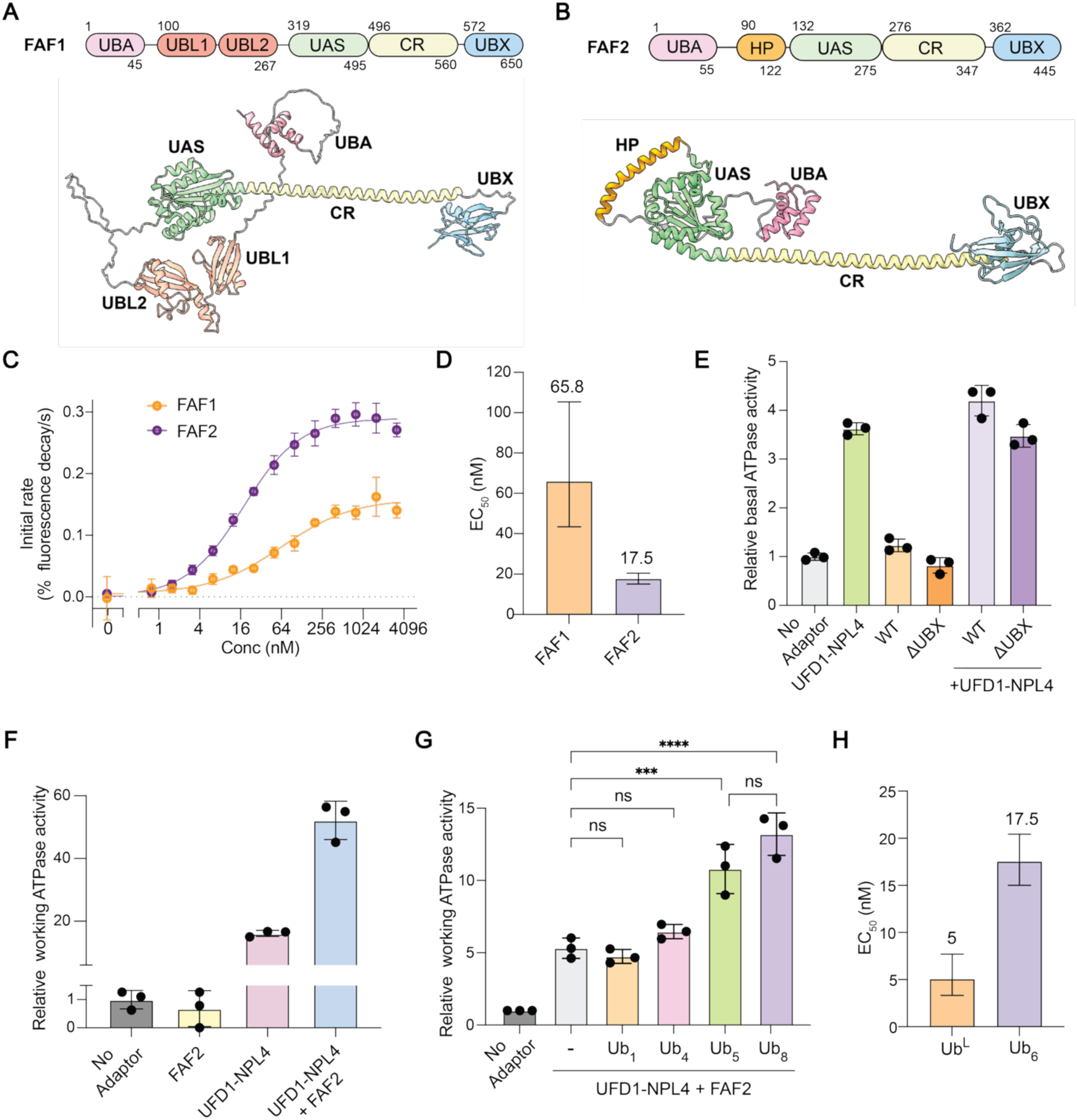
Enhanced unfolding of ubiquitylated substrate by FAF2. (A) Domain organization (top) and Alphafold3 structure (bottom) of full-length FAF1, highlighting the UBA, UBL1, UBL2, UAS, CR, and UBX domains. (B) Domain organization (top) and Alphafold3 structure (bottom) of full-length FAF2, highlighting the UBA, Hairpin (HP) motif, UAS, CR, and UBX domains. (C) Initial unfolding rate (first 2 min) of unfolding of Ub_6_-Eos as a function of FAF1 and FAF2 concentration. p97 (400 nM) and UN (400 nM) concentrations were kept constant. Data shown as mean ± SD for n = 4 technical replicates. (D) Comparison of EC_50_ values of FAF1 and FAF2 for enhancing the unfolding rate of Ub_6_-Eos substrate by p97-UFD1-NPL4 complex. EC_50_ values were calculated from data in (C). Error bars denote 95% CI. (E) Relative ATPase activity of p97 without or with the indicated cofactors. Basal ATP hydrolysis was measured and normalized to p97 alone (no adaptor). Data represent mean ± SD for n=3 technical replicates. (F) Relative ATPase activity of p97 with indicated cofactors in the presence or absence of Ub^L^ substrate. Working ATP hydrolysis was measured and normalized to p97 alone. Data represents mean ± SD for n=3 technical replicates. (G) Relative ATPase activity of p97 with indicated cofactors in the presence of monoubiquitin and free K48-linked polyUb chains of various lengths. Working ATP hydrolysis was measured and normalized to p97 alone. Data represents mean ± SD for n=3 independent experiments, each with 3 technical replicates. (H) Comparison of EC_50_ values of FAF2 for enhancing the unfolding rate Ub^L^ and Ub_6_-Eos substrate by p97-UFD1-NPL4 complex. EC_50_ values were calculated from data shown in Figure 2C and EV7C. Error bars denote 95% CI. Statistical significance was calculated by one-way ANOVA followed by multiple-comparison testing (ns, not significant P > 0.05; *P ≤ 0.05; **P ≤ 0.01, ***P ≤ 0.001, ****P ≤ 0.0001).

Using K48-linked Ub_6_-Eos as a model substrate, we found that stoichiometric amounts of either adaptor (one FAF1 or FAF2 per p97–UN complex) was sufficient to achieve maximal unfolding rates, with no further enhancement at higher FAF1/FAF2: p97-UN ratios (**Figure 2C**). Notably, FAF2 showed substantially higher maximal unfolding rate and displayed a lower EC_50_ (17.5 nM) compared to FAF1 (65.9 nM) (**Figure 2C, D**). To understand the determinants underlying this effect, we tested whether FAF2 enhances unfolding by modulating ATP hydrolysis activity of p97. We hypothesized of FAF2 binding to the p97–UN complex might stabilize a conformational state of p97 that facilitates rapid nucleotide turnover and ATP-dependent substrate unfolding. To test this possibility, we measured the basal ATPase activity of p97 by an endpoint phosphate release assay. While UN significantly enhanced the basal ATPase activity of p97, the addition of FAF2 to p97 by itself or to p97-UN complex did not further increase ATP hydrolysis (**Figure 2E**), indicating that the FAF2-mediated enhancement of p97 activity is dependent on the presence of ubiquitylated client. Consistent with this, FAF2 markedly increased ATP hydrolysis by the substrate-engaged p97–UN complex (**Figure 2F**), demonstrating that FAF2 affects substrate-stimulated rather than basal ATPase activity.

Having identified activation to be polyUb-dependent, we next asked how ubiquitin chain length influences this stimulation and if there is a chain length threshold for FAF2-mediated activation of the p97–UN complex. Hence, we measured phosphate release upon addition of free K48-linked Ub chains of varying lengths to the p97–UN– FAF2 complex (**Figure EV2A**). Mono-Ub and tetra-Ub (Ub_4_) substrates did not increase ATP hydrolysis, whereas a penta-Ub (Ub_5_) chain significantly enhanced ATP hydrolysis. Extending the chain to octa-Ub (Ub_8_) only caused a minor increase in phosphate release over what was observed for Ub_5_, indicating that a penta-Ub chain is sufficient for stimulation of ATPase activity (**Figure 2G**). In line with this observation, FAF2 promoted unfolding of both short (Ub_6_-Eos) and long (Ub^L^-Eos) chain modified substrates, albeit with a modest difference in potency, displaying a lower EC_50_ for the longer chain (5 nM) compared to the shorter chain (17.5 nM) (**Figure 2H**). Collectively, FAF2 enhances p97-UN-mediated substrate unfolding and ATP hydrolysis once a threshold ubiquitin chain length of at least 5 K48-linked ubiquitins is reached.

### FAF2 forms a stable complex with p97 in a UFD1-dependent manner

To uncover how FAF2 forms a complex with p97, we used AlphaFold3 to model the p97–UN–FAF2 complex, incorporating the p97 N–D1 hexamer, UFD1, NPL4, and FAF2. All predicted models show a consistent interaction pattern in which three p97 protomers engage adaptors bound to the N domain through the UBX-L domain of NPL4, the UBX domain of FAF2, and the SHP1 and SHP2 motifs of UFD1 (**Figure 3A, EV2B**). Of note, the FAF2 UBX domain is predicted to bind the N domain of the same p97 protomer where the SHP1 motif of UFD1 engages, whereas the SHP2 motif of UFD1 interacts with the adjacent p97 protomer. The N domains are modelled in the ‘up’ conformation, resembling the state when ATP is bound to both the D1 and D2 domains. The models position the NPL4 tower on top of the p97 hexamer, with UFD1 located adjacent to it through interactions with the MPN domain of NPL4. FAF2 is positioned on the same side of the hexamer, where the CR bridges the p97 N domain to the UFD1–NPL4 complex (**Figure 3A**). Notably, the CR contacts the UT3 region of UFD1 but makes no direct contact with NPL4. The UBA, HP, and UAS domains of FAF2 do not engage the p97–UN complex in the absence of substrate (**Figure 3B**).

**Figure 3:**
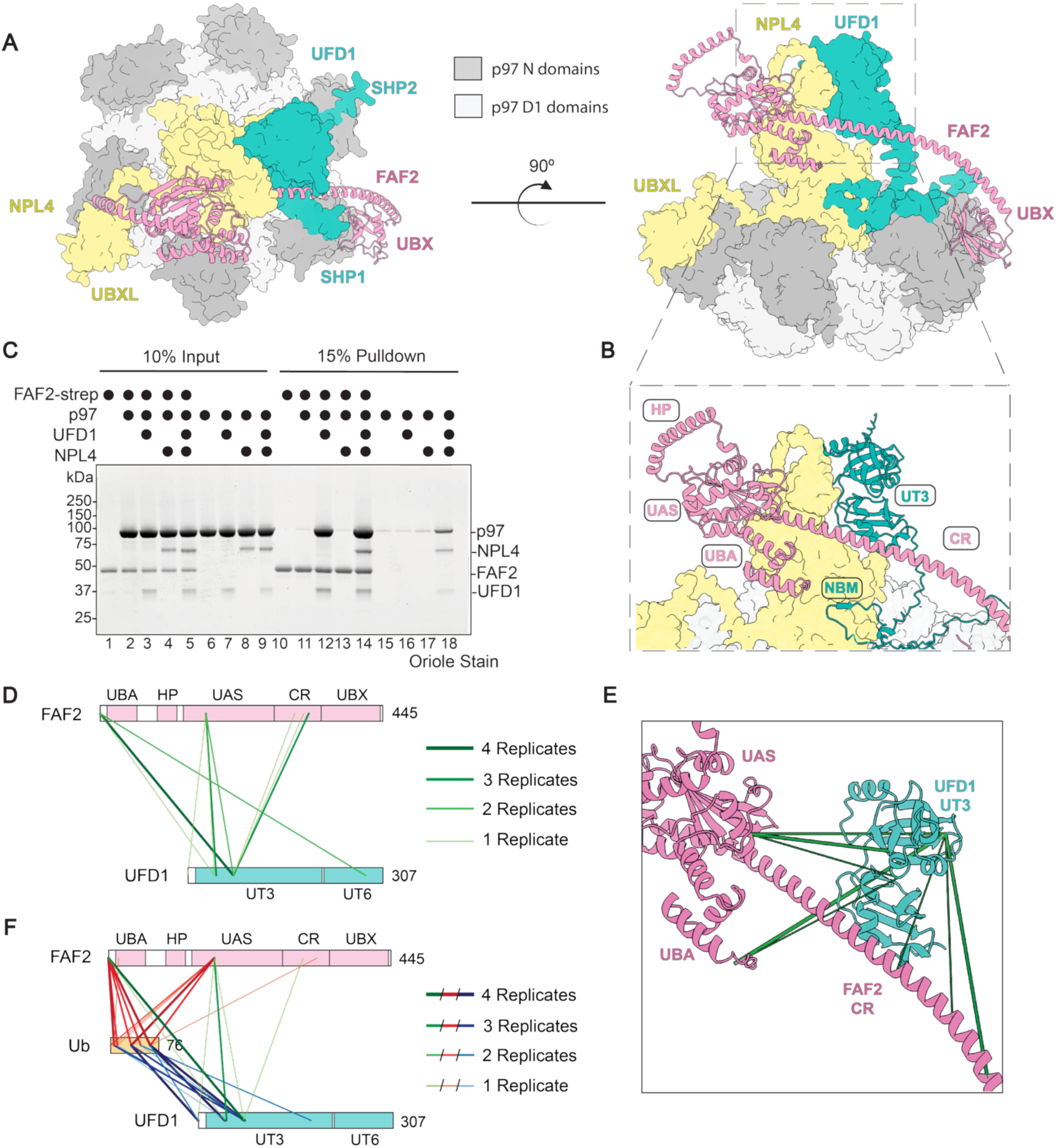
FAF2 engages the p97–UFD1–NPL4 complex through multivalent interactions. (A) AlphaFold3-predicted structure of the p9-ND1 hexamer bound to the UFD1–NPL4 heterodimer and FAF2. p97 N domains are shown in dark grey and D1 domains in light grey; UFD1 is shown in teal, NPL4 in yellow, and FAF2 in pink. Left, top view of the complex highlighting occupancy of p97 N domains by UBX domains and SHP motifs of cofactor proteins. Right, side view of the complex illustrating the extended architecture of FAF2 bridging p97 N domain and the UFD1–NPL4 complex. (B) Close-up view of the FAF2 C-terminal region bound to UFD1–NPL4, highlighting interactions mediated by the CR of FAF2 and the UT3 domain of UFD1, while UT6 domain interacts with NPL4. (C) Strep pull-down of C-ter Twin-Strep-tagged FAF2 performed with p97, individual UFD1 or NPL4, the UFD1–NPL4 complex, or the p97 and UFD1–NPL4 complex together, followed by Oriole staining, showing that FAF2 binding requires both p97 and UFD1. (D) Crosslinking of p97: UFD1-NPL4: FAF2 ternary complex with DSSO crosslinker. All detected crosslinks between the components are shown for n=4 replicates. The colour and width of the crosslinks indicate the reproducibility across different replicates. (E) Crosslinks from (D) were mapped onto the AlphaFold3 model of p97: UFD1-NPL4: FAF2 ternary complex. The FAF2 and UFD1 structures are highlighted to illustrate the crosslinks between them. (F) Crosslinking of p97: UFD1-NPL4: FAF2 ternary complex in presence of polyUb substrate (free Ub_6_ – Ub_8_ chains) with DSSO crosslinker. All detected crosslinks between the components are shown for n=4 replicates. The colour and width of the crosslinks indicate the reproducibility across different replicates.

To experimentally test the verity of the prediction, we performed *in vitro* pulldown assays using Twin-Strep–tagged FAF2 and assayed binding to p97 by itself or in the presence of additional components. This revealed that FAF2 pulled down only minimal amounts of p97 when incubated alone or with NPL4 (**Figure 3C**, lanes 11 and 13). In contrast, the presence of UFD1 markedly increased p97 binding to FAF2, whereas the presence of UN complex did not further enhance the interaction (**Figure 3C**, lanes 12 and 14). These results indicate that UFD1 is both necessary and sufficient to mediate FAF2 recruitment to the p97 complex, consistent with the CR of FAF2 engaging the UT3 region of UFD1 in the structural model. Importantly, under the same conditions, no interaction between FAF2 and UFD1 was detected in the absence of p97, as the pulldown of FAF2 was unable to co-purify UFD1 when p97 was not added (**Figure EV2F**), suggesting that a multivalent interaction network stabilizes FAF2 binding to the assembled p97-UN complex. These results were further confirmed by mass photometry, which revealed the formation of a 1:1:1 complex of hexameric p97 with UFD1, NPL4 and FAF2 (**Figure EV2C**).

To get more insights into the molecular interactions that enable complex formation between FAF2 and p97-UN, we analysed p97-UN-FAF2 complexes by crosslinking mass spectrometry (XL-MS). In line with structural models, multiple crosslinks are detected between the N domain of p97 and all the other components, including crosslinks to the UT6 domain of UFD1 and the UBX-L and ZF-Npl4 domains of NPL4 (**Figure 3D, EV3**). The D1 domain of p97 also makes contacts with the UT6 domain of UFD1 and the MPN domain of NPL4. The UT6 domain of UFD1 makes multiple crosslinks to the MPN domain of NPL4. The UT3 domain of UFD1 makes interactions with the MPN domain of NPL4 and the UAS domain of FAF2. Crosslinks are detected between the CR of FAF2 and the UT3 domain of UFD1 primarily in the absence of ubiquitin but seem to be reduced in the presence of ubiquitin, suggesting conformational rearrangements upon substrate engagement. Of note, very few peptides were detected in the CR region, likely due to the amino acid composition in this stretch, which limits resolution in this region. The crosslinks between the rest of the components do not appear to alter significantly in the absence or presence of ubiquitin. When present, ubiquitin forms extensive crosslinks with the MPN domain of NPL4, the UT3 domain of UFD1, and the UAS, and CR of FAF2, suggesting it engages multiple domains of adaptors to coordinate substrate engagement (**Figure 3F, EV3**).

Hydrogen deuterium exchange (HDX-MS) analysis uncovered additional insights into the dynamic behaviour of the p97-UN-FAF2 complex. We generated a comprehensive peptide map of the p97–UN complex and assessed how FAF2 binding alters solvent exchange across individual components of the complex. Notably, FAF2 association resulted in a pronounced reduction in solvent exchange within the UT3 domain of UFD1 (residues 133–149), consistent with direct engagement of FAF2 at this site and in strong agreement with the AF3 prediction (**Figure 4A, 4B**). Interestingly, FAF2 binding also led to reduced solvent exchange within the UFD1 SHP2 motif (**Figure 4C, 4D**). AF3 predicts that UFD1-SHP1 and FAF2-UBX bind distinct sites on the same p97 N domain, whereas the SHP2 motif interacts with a neighbouring N domain. The reduced solvent exchange observed in this region suggests that either it engages the same p97 N domain as FAF2 or FAF2 binding induces conformational rearrangements to stabilize UFD1 on p97 via the SHP2 motif. Similarly, corresponding regions of the p97 N domain, where both the SHP2 motif of UFD1 and the UBX domain of FAF2 bind, also exhibited reduction in solvent exchange. In addition, regions in the CTD of NPL4 also showed minor protection from solvent exchange in presence of FAF2, indicating potential allosteric effects (**Figure EV2G**). One limitation of this analysis is that changes in p97-derived peptides occurring on one protomer of p97 were diluted by six-fold due to stoichiometric effects, likely underestimating the effect of FAF2 binding on p97 conformation.

**Figure 4:**
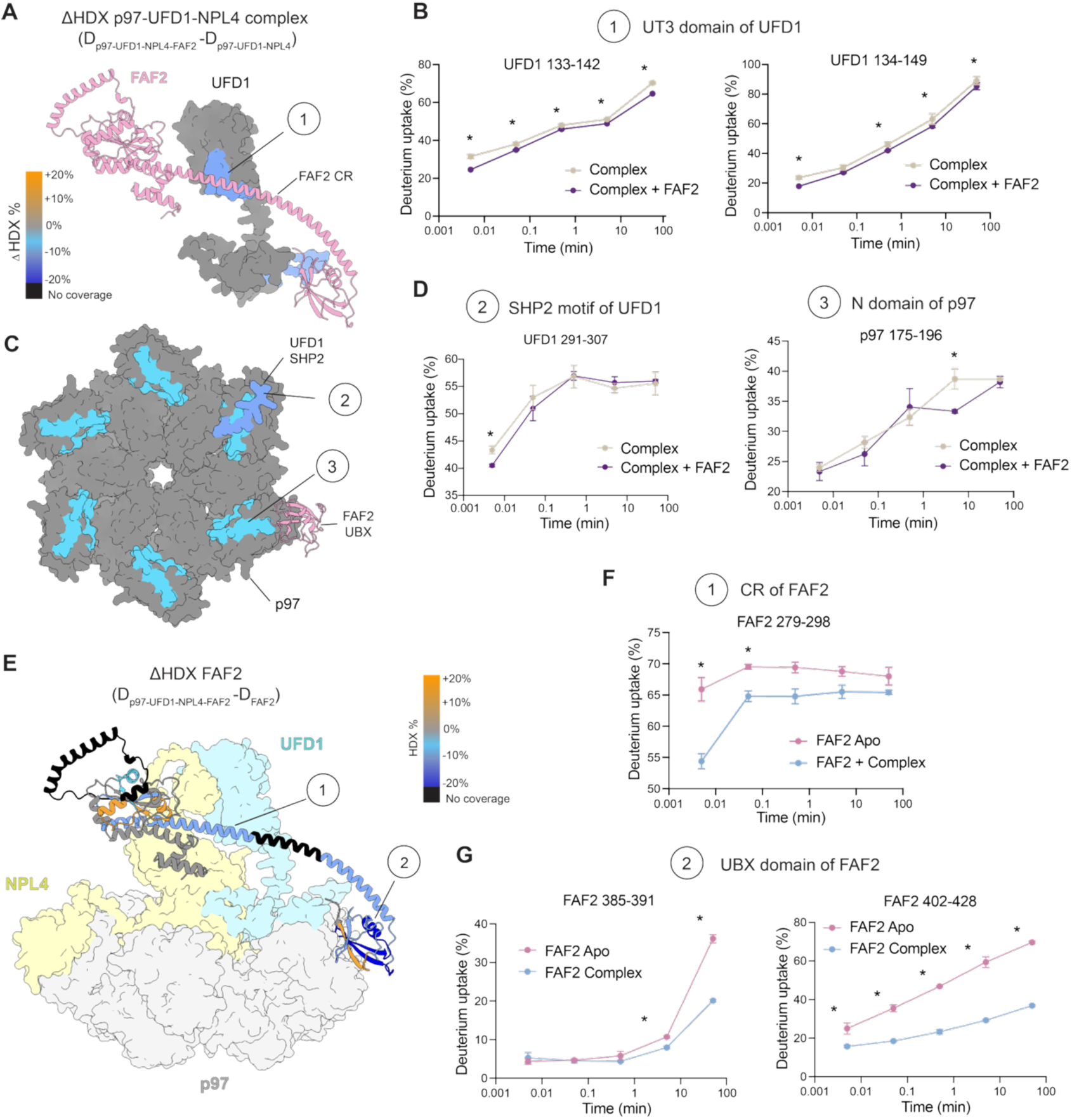
Hydrogen–deuterium exchange (HDX-MS) analysis of FAF2 interactions with the p97–UFD1–NPL4 complex. (A) HDX-MS was performed with p97:UFD1-NPL4:FAF2 complex in the absence or presence of FAF2 at different time points. Regions in UFD1 which exhibited a reduction of solvent exchange upon FAF2 binding are highlighted. Regions with reduced solvent exchange are shown in blue, increased exchange in orange, no coverage in black, and no significant change (−5% to +5%) in grey. (B) Deuterium uptake kinetics of selected peptides in the UFD1 UT3 domain showing FAF2-dependent changes. Data represent mean ± SD (n = 3 independent replicates). (C) ΔHDX mapping of p97 and the UFD1 C-terminus upon FAF2 binding using the same colour scheme as in (A). (D) Deuterium uptake kinetics of selected peptides within the UFD1 SHP2 motif and corresponding binding sites in the p97 N domains. Data represented as mean ± SD. (E) HDX-MS analysis of FAF2 in the apo state or bound to the p97:UFD1:NPL4 complex, displayed using the colour scheme described in (A). (F-G) Differential deuterium uptake of selected FAF2 peptides in the CR and UBX domain upon complex formation. Data represent mean ± SD (n = 3 independent replicates). * = > 5% and >0.5 Da difference and passes a student t-test with p = 0.05 (or lower).

Notably, HDX–MS analysis also revealed a key aspect of FAF2 secondary structure. While AF3 predicted the central region in between UAS and UBX domain of FAF2 (residues 275–348) to be a long α-helix, HDX-MS analysis of FAF2 in apo state shows a very rapid solvent exchange for peptides in this region even at very early time points (0.3 s), a feature typical of an intrinsically disordered region (**Figure 4E**). Remarkably, when FAF2 binds the p97–UN complex, deuterium uptake in the peptides of this region decreases markedly, indicating that this region adopts a more ordered secondary structure, likely consistent with the α-helical structure predicted by AF (**Figure 4E, 4F**). Additionally, peptides in the FAF2-CR, which binds UT3 of UFD1, and FAF2-UBX, which engages the p97 N domain, showed decreased solvent exchange, further validating the structural model. However, amino acid composition within the UFD1 binding interface did not allow sufficient coverage of the entire region. Nevertheless, the differential solvent exchange properties clearly demonstrate how FAF2 induces coordinated stabilisation across UFD1 and p97 N domains through multivalent interactions, providing mechanistic insights into how FAF2 stimulates the activity of the p97-UN complex.

### FAF2 activation motif bridges UFD1-ubiquitin interactions to drive p97 activation

To gain insights into how FAF2 activates the unfolding of substrates carrying both short and long Ub chains, we performed AF3 predictions of p97-UN-FAF2 complex together with 10 ubiquitin moieties (**Figure 5A, EV4A**). This revealed multisite interactions between polyUb and the p97 complex. In line with our XL-MS results (**Figure EV2D, EV3**), ubiquitin interactions with the MPN and CTD domain of NPL4, the UBA domain and CR of FAF2 and the UT3 domain of UFD1 are consistently recapitulated in four out of the five models generated (**Figure EV5**). Intriguingly, two ubiquitin moieties simultaneously engaged by UFD1 UT3 domain are positioned such that the C-terminus of the substrate-proximal ubiquitin points towards K48 of the adjacent distal ubiquitin molecule (**Figure 5B**). The UT3 domain interacts with the Ile44 patch of the proximal Ub in a similar manner as characterized for yeast Ufd1 (Williams *et al*, 2023) (**Figure 5D**). The distal Ub binds the entrance of the NPL4 groove roughly where the initiator ubiquitin would be positioned and inserted, as observed in the yeast complex (Twomey *et al*, 2019). Importantly, the FAF2-CR sits atop this UFD1 UT3–diUb complex, making contacts mostly with the Asp58 patch of the proximal Ub and the UT3 domain (**Figure 5C, 5E**). The FAF2-CR also contacts the K48-linkage between the proximal and distal Ub (**Figure 5B, 5C**), thus likely stabilizing the UT3-diUb interactions. Based on the structural model and the observation that FAF2 reduces the Ub chain length threshold for substrate unfolding, we predict that ubiquitin interactions with the FAF2 UBA domain and/or the CR would be important for this function. To map the region of FAF2 critical for stimulating p97 activity, we tested three key truncations of FAF2 – ΔUBA (1-137), ΔCR (296-349) and ΔUBX (362-445) in an unfolding assay (**Figure 5F**). Deletion of the UBX domain completely abolished unfolding activity, confirming that engagement with p97 via the UBX domain is essential for its function. In contrast, removal of the UBA domain had no effect on the rate and efficiency of substrate processing indicating that direct ubiquitin binding via this domain to FAF2 is not required for its stimulatory role. Importantly, deletion of a part of the CR also fully abolished substrate unfolding activity, indicating that this region is indispensable along with the UBX domain for FAF2 function (**Figure 5F**). Furthermore, a similar domain dependence was also reflected in substrate-induced ATP hydrolysis by the p97–UN–FAF2 complex. Deletion of either the UBX or CR strongly reduced phosphate release, whereas removal of the UBA domain showed only a modest loss in activity **(Figure 5G**).

**Figure 5:**
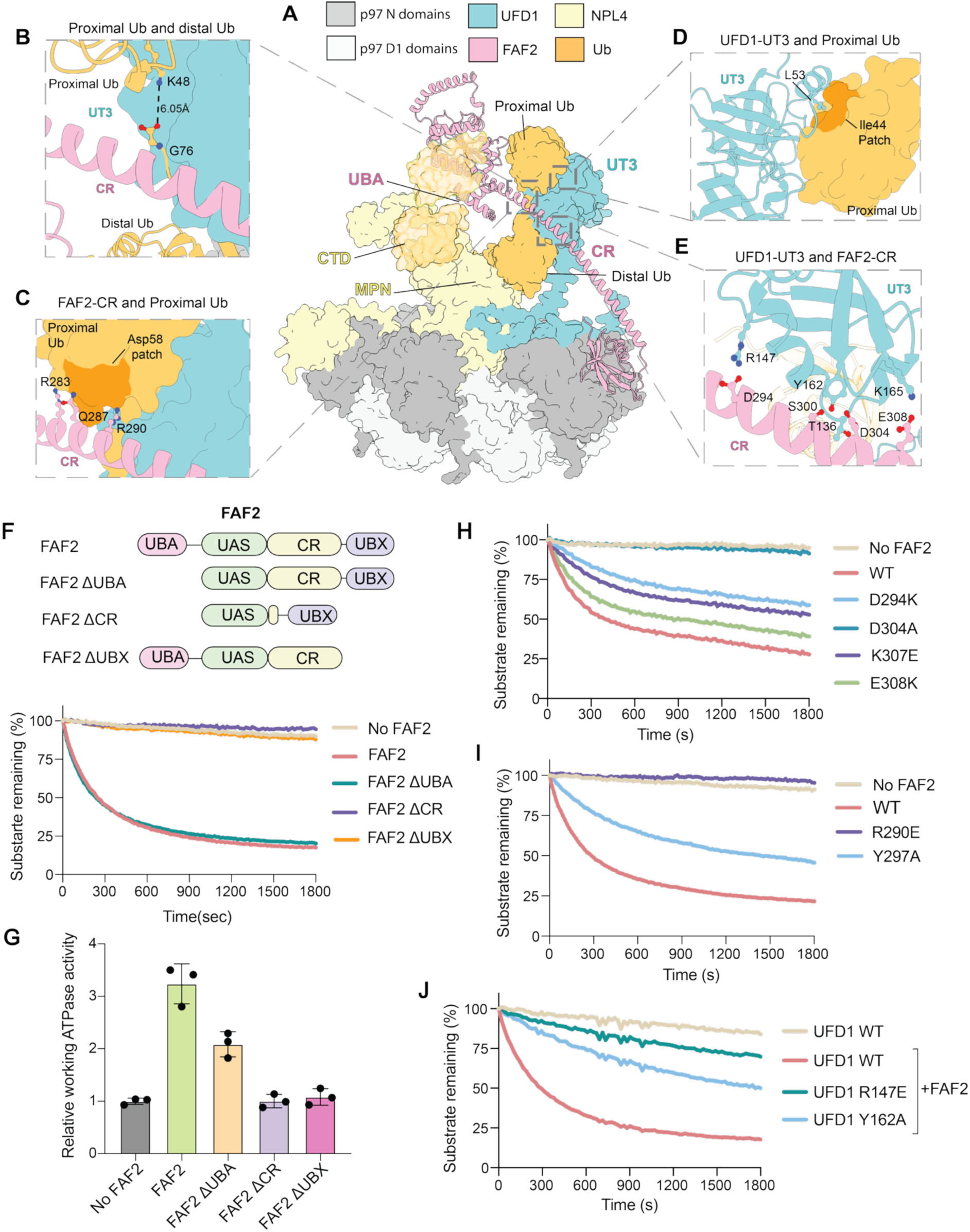
Coordinated ubiquitin engagement by FAF2 and UFD1 enables p97-mediated substrate unfolding. (A) Alphafold3 model of p97 N-D1 hexamer, UFD1-NPL4, FAF2 and 10 ubiquitin molecules. The model shows binding of 4 ubiquitin across various ubiquitin-binding domains of FAF2, UFD1 and NPL4. (B) Inset showing two ubiquitin molecules bound to UFD1 and FAF2 in a K48-linked conformation. (C) Enlarged view of the FAF2-proximal ubiquitin, illustrating the FAF2-CR–Asp58 patch interaction. (D) The UFD1-proximal Ub interface shows engagement with the Ile44 patch of ubiquitin. (E) UFD1 UT3 and FAF2 CR interface remain mostly unaltered in presence of ubiquitin. The residues mediating interactions are shown as ball and stick. (F) Top: Schematic representation of FAF2 truncation constructs used. Bottom: Unfolding of Ub_6_–Eos by p97–UFD1–NPL4 in the presence of WT FAF2 or FAF2 truncation variants. (G) ATPase activity of p97 in the presence of the indicated FAF2 truncations and Ub^L^ substrate, normalized to ATPase activity of p97 in the presence of the UFD1–NPL4 complex and substrate. Data represent mean ± SD (n = 3 technical replicates) and extended from the data shown in Figure 2F. (H-I) Analysis of unfolding activity of FAF2 mutants predicted by Alphafold3 to disrupt UFD1 binding, measured in presence of UFD1-NPL4. (J) Unfolding activity of UFD1 mutants predicted by Alphafold3 to impair interaction with FAF2 binding, measured in presence of FAF2. For (F, H-J) traces were normalized to fluorescence intensity at time 0 and substrate-only controls. Data is shown for n = 4 technical replicates.

FAF2 and UFD1 UT3 domain interactions are mediated primarily by polar electrostatic and hydrogen bonding interactions between D294, Y297, S300, D304, K307 and E308 in FAF2 and R147, T136, Y162, K165 in UFD1, as suggested by the AF3 model and supported by the HDX-MS analysis (**Figure 5E**). These interactions are also largely conserved in the FAF1-p97-UN complex, suggesting a conserved mode of interaction between FAF1 and FAF2 (**Figure EV5**). To disrupt the FAF2 CR–UFD1 UT3 interaction, we introduced point mutations in FAF2 and UFD1 guided by conservation analysis across 150 eukaryotic species (**Figure EV2E**). Among the mutations tested, the D304A mutant resulted in a complete loss of the unfolding activity, whereas D294E, Y297A, K307E and E308K partially reduced unfolding (**Figure 5H, 5I, EV4C, EV4D**). Complementary mutations on UFD1 UT3 domain (R147E and Y162A), which interact with D294 and E304 of FAF2, similarly impaired unfolding activation (**Figure 5J, EV4E**). Consistent with these functional defects, the FAF2 D304A mutant markedly reduced binding to the p97–UN complex (**Figure EV4G**). Likewise, mutation of R147 and Y162, the residues in UFD1 that mediate contacts with D294 and D304 of FAF2, disrupted FAF2 binding (**Figure EV4H**).

We also mutated the residues within the FAF2 CR mediating interactions with ubiquitin to probe the importance of ubiquitin interactions. Mutation of FAF2 R290, which contacts the K48 linkage in the AF3 model, fully abolished the unfolding stimulation, whereas others showed no significant effect (**Figure 5I, EV4B, EV4C**). Unexpectedly, the R290E mutation also abrogates interaction of FAF2 with p97-UFD1-NPL4 complex in the absence of ubiquitin (**Figure EV4G**). This finding raises the possibility that R290 in FAF2 adopts a slightly different conformation upon ubiquitin binding, enabling simultaneous recognition of UFD1 and ubiquitin by the UT3 domain. Mutation of UFD1 L53, which engages the proximal ubiquitin in our model, strongly reduced the unfolding activity, supporting the predicted positioning of ubiquitin in the complex (**Figure 5D; Figure EV4F**). Collectively, these observations demonstrate that the CR-mediated FAF2–UFD1 interactions, together with FAF2 engagement of proximal ubiquitin, are essential for stabilizing productive p97 assemblies and promoting efficient substrate unfolding.

Although FAF2-dependent activation required the presence of ubiquitin, deletion of the UBA domain, the only characterized ubiquitin-binding region within FAF2, had no effect on either unfolding or ATP hydrolysis. We therefore asked whether the FAF2 CR, which is essential for its activity, might contribute to ubiquitin binding. To address this, we performed pulldown assays using Strep-tagged-FAF2 (ΔUBA) in the presence of Ub trimer (Ub_3_) and hexamer (Ub_6_), with full-length FAF2 included as a positive control. The results showed that full-length FAF2 selectively pulled down ubiquitin hexamers but not trimers, indicating that the UBA domain in FAF2 preferentially binds longer ubiquitin chains. In contrast, the FAF2 ΔUBA construct did not show any detectable binding to either Ub3 or Ub6, confirming that the CR of FAF2 is not a ubiquitin-binding domain (UBD) (**Figure EV4I**). These results support the structural model wherein the FAF2-CR by itself does not have high enough affinity for ubiquitin binding but bridges UFD1 UT3 and proximal Ub by multivalent interactions to form a high-avidity complex. In support of this cooperative model, we were unable to detect binding of either FAF2 or UFD1 to K48-linked diUb chains by isothermal titration calorimetry (ITC) (**Figure EV4J, EV4K**). Based on our structural and biochemical data, we identify residues 286–312 of FAF2 as the minimal UFD1-interacting region required for unfolding stimulation and refer to this segment as the activation motif (AM). Together, these results support a cooperative binding model in which the FAF2 activation motif enhances UFD1’s effective affinity for K48-linked ubiquitin chains through multivalent interactions. This cooperative mechanism provides a molecular explanation for how FAF2 reduces the ubiquitin chain length threshold required by promoting productive p97-UN substrate engagement.

### FAF2 activation motif scaffolding guides de novo design of p97 activator

Loss-of-function variants of p97, such as D395G and D592N, show reduced ATPase activity and defects in clearance of aggregates (Ju *et al*, 2008; Weihl *et al*, 2006). These observations suggest that activating p97 could facilitate the clearance of aggregation-prone proteins in cells, offering a potential therapeutic strategy for proteinopathies. Building on the principle that the FAF2 AM is essential for UFD1 binding and accelerating the unfolding reaction, we explored the idea of designing activators of the p97-UN complex through a *de novo* protein design approach.

To this end, we used RFdiffusion (Watson *et al*, 2023) to scaffold potential p97-activating proteins around the FAF2 activation motif (286-312). The motif scaffolding was performed in the presence of the UT3 domain of UFD1 (17-196) and two ubiquitin molecules positioned in K48-linked orientation obtained from the AF3 model of the complex as described before (**Figure 6A**). This structural model served as the input for binder generation using RFdiffusion (Watson *et al*, 2023), integrated with ProteinMPNN for amino acid sequence design (Dauparas *et al*, 2022) and AlphaFold2-based structure prediction and validation **(Figure EV6A**). Residues in the FAF2 AM involved in UFD1 and ubiquitin interactions (T286, L289, R290, Q292, Q293, D294, A296, Y297, S300, L301, D304, Q305, K307, E308, and K311), as predicted by the AF3 model, were held fixed during design. Various N- and C-terminal extensions were then sampled, while RFdiffusion redesigned the intervening residues within the FAF2 AM helix, optimizing the sequence to pack efficiently against the fixed interaction surfaces (Wang *et al*, 2022). This approach maintained the FAF2 AM in the three-dimensional conformation it adopts upon UFD1 binding, thereby preserving the interaction surfaces with UFD1 and ubiquitin, while allowing the surrounding scaffold to reorganize and stabilize around it. The design process generated a diverse set of candidate scaffolds, representing multiple distinct backbone trajectories. Candidates were filtered for structural confidence, resulting in a pool of binder designs that preserve the geometry required for UFD1 and ubiquitin engagement (**Supplementary Table S2**). In total, 15 binder designs were selected for experimental validation.

**Figure 6:**
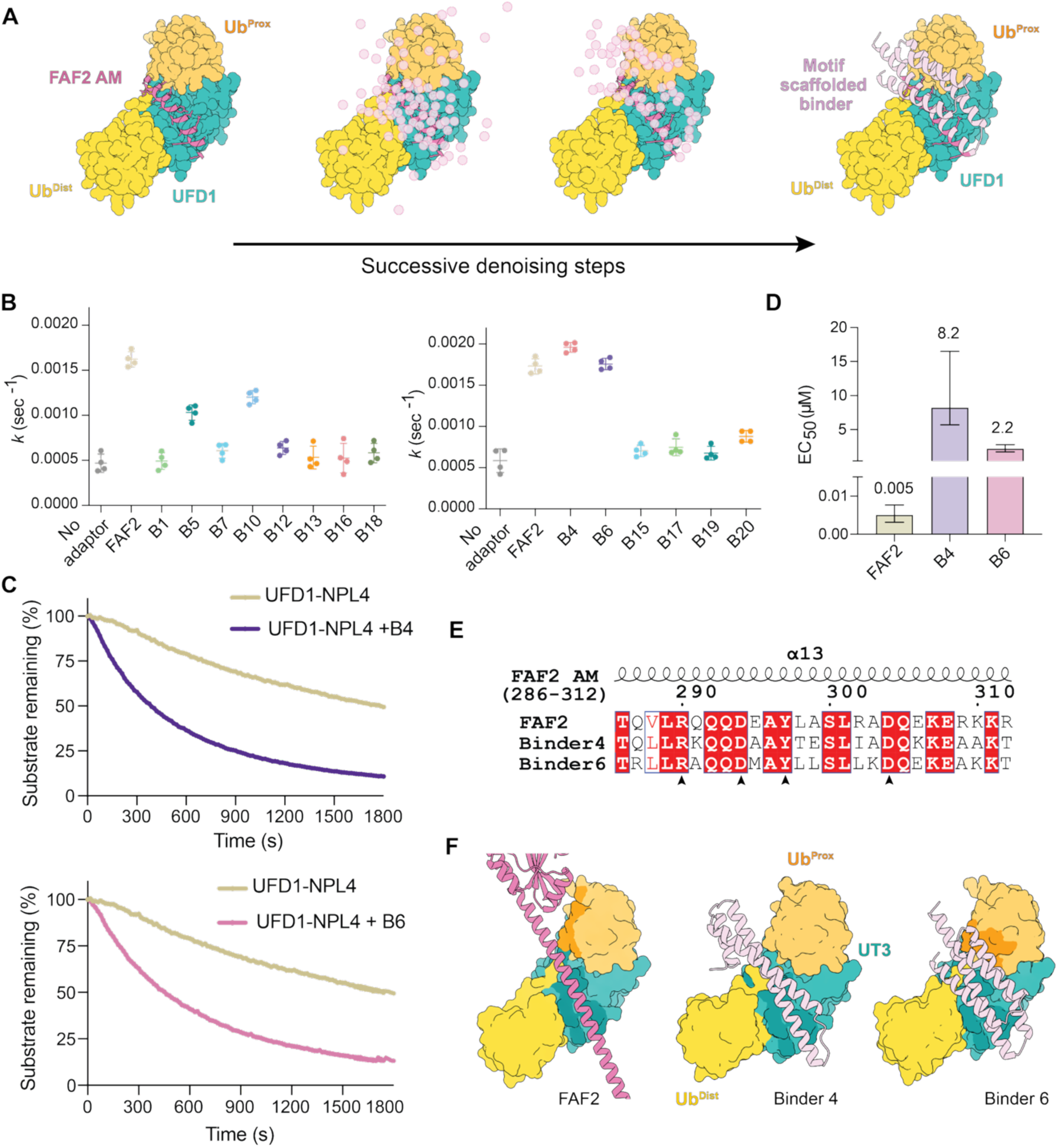
Targeting UFD1 with de novo binders enhance p97-mediated substrate unfolding activity. (A) De novo UFD1 binders were designed by fixing residues within the FAF2 motif (residues 286–312, dark pink) in the UFD1 (teal)–di-ubiquitin (orange) complex and building a scaffold around it (light pink) via RFdiffusion, transforming noisy residue frames into a stable backbone engaging the target. (B) Screening of FAF2-motif–scaffolded binders for enhancement of p97-mediated substrate unfolding in a UFD1-NPL4-dependent manner. Unfolding of a Ub^L^ substrate were measured in the presence of p97–UFD1–NPL4 and individual binders, with FAF2 as a positive reference. Rate was calculated by single exponential fitting of the trace. Data represent mean ± SD from n = 4 technical replicates. (C) Single turnover unfolding traces of Ub^L^ in the absence or presence of representative binders show potent activation of unfoldase complex. Data represent mean ± SD from n = 4 technical replicates. (D) Comparison of EC₅₀ values for enhancement of p97-mediated substrate unfolding by FAF2 and selected scaffolded binders, calculated from plotting the unfolding rates from data in Figure EV7 as a function of FAF2 or binder concentration. Data represent mean ± SD from n = 3 technical replicates for FAF2 and n=4 technical replicates for binders. Error bars denote 95% CI. (E) Sequence alignment of the functional FAF2 motif and selected scaffolded binders highlighting conserved residues that were kept fixed during RFdiffusion-based binder design. FAF2 numbering and secondary structure elements are shown. Black arrows indicate FAF2 residues required for UFD1 binding and enhanced unfolding activity, as suggested by the structural model and validated by mutational analysis. (F) AlphaFold models of FAF2 and engineered binders bound to the UFD1 UT3 and diUb complex. FAF2 is shown in dark pink, UT3 in teal, proximal Ub in orange and distal Ub in yellow. Binder 4 and Binder 6 engage the complex in a similar overall orientation as FAF2 and contact largely overlapping surfaces on UT3 and ubiquitin.

Of the 15 binders tested, 14 expressed well as soluble proteins (**Figure EV7A)**. We next assessed their ability to stimulate p97 activity by monitoring unfolding of Ub^L^ substrate. Remarkably, among the 14 binders tested, 4 binders (Binder 5, Binder 10, Binder 4 and Binder 6) enhanced substrate unfolding in presence of UFD1-NPL4 under the same experimental conditions as FAF2 (**Figure 6B, EV7B**). The extent of stimulation varied across designs, with Binder 4 and Binder 6 showing the strongest effect with similar unfolding kinetics as observed with FAF2 **(Figure 6B, 6C**). However, both binders exhibited ∼500-1000-fold higher EC₅₀ values (Binder 4 EC_50_ = 8.2 µM, Binder 6 EC_50_ = 2.2 µM) compared with FAF2 (EC_50_ = 6 nM), indicating reduced potency despite achieving a similar maximal unfolding activity **(Figure 6D, Figure EV7C)**. Sequence alignment of the *de novo* activators (Binder 4 and Binder 6) with FAF2 revealed strong conservation of residues within the FAF2 helical activation motif, whereas intervening residues showed great variability **(Figure 6E),** indicating that activity is primarily driven by the key interaction surface. Structural analysis of binder 4 and binder 6 in complex with UT3–diUb, and comparison with the corresponding FAF2-bound model, revealed that these de novo binders retain interactions with the UT3 domain as observed for FAF2. While binder 4 shows no additional contacts with ubiquitin, binder 6 engages both ubiquitin moieties as well as UFD1 (**Figure 6F, EV7D**), explaining lower EC_50_ for binder 6. Together, these results demonstrate that the key UFD1-UT3 and ubiquitin-contacting residues within the FAF2 AM are both necessary and sufficient to confer activation of p97-UN complex.

## Discussion

The core function of p97, the ATP-driven mechanical unfolding of ubiquitylated proteins, is highly conserved from yeast to human. However, human p97 has evolved into a more tightly regulated ATPase relying on cofactors that tailor p97 activity in a substrate- and context -dependent manner, which is necessary given that p97 must unfold diverse and challenging substrates across multiple subcellular compartments. This correlates with an expansion in the number of cofactors, wherein individual cofactors functionalize p97 to perform specialized functions in distinct cellular pathways including selective organellar quality control such as FAF2-driven pexophagy and mitophagy (Koyano *et al*, 2024; Zheng *et al*, 2022; Ganji *et al*, 2023), lysophagy mediated by UBXD1 and PLAA (Papadopoulos *et al*, 2017) and the disassembly of aggresomes and stress granules by FAF1 and FAF2 (Körner *et al*, 2025; Turakhiya *et al*, 2018; Gwon *et al*, 2021).

Despite detailed biochemical and structural characterization of the yeast Cdc48-UN complex, our study and previous work reveal surprising differences to the human system (Fujisawa *et al*, 2022). Human p97-UN, for unknown reasons, displays substantially lower unfolding efficiency *in vitro* compared to yeast, even when unfolding substrates modified with long ubiquitin chains. One possible explanation is that yeast Ufd1-Npl4 binds polyubiquitin with higher affinity, allowing more efficient substrate engagement. This deficit, we believe, is compensated in humans by activating cofactors. Our finding that FAF1 and FAF2 act in concert with UFD1-NPL4 and accelerate the activity of human p97 supports this idea. Biochemical analysis of the FAF2–p97 complex revealed that FAF2 binding to p97 is highly dependent on the presence of UN, contrasting with studies on the homologue FAF1 that reported direct FAF1–p97 interaction (Ewens *et al*, 2014). In addition, our screen revealed p47 and UBXN7 to potentiate unfolding of substrates modified with long polyUb chains in an UFD1-NPL4-dependent manner. This is consistent with previous reports implicating UBXN7 and FAF1 in mammalian replisome complex disassembly (Fujisawa *et al*, 2022). p47 is unique amongst these activating cofactors in that it contains an additional SEP domain, which is also present in adaptors such as UBXN2B (p37) and UBXN2A (p47L). While UBXN2A and 2B unfold substrates in a ubiquitin-independent manner (Kracht *et al*, 2020; Weith *et al*, 2018), p47 also contains a UBA domain and does not support this ubiquitin independent activity (Kracht *et al*, 2020). Intriguingly, we here found that p47 partners with UFD1-NPL4 to stimulate ubiquitin dependent unfolding. This observation challenges the prevailing model that these adaptors bind p97 in a mutually exclusive manner (Meyer *et al*, 2002). However, the underlying mechanism for this cooperativity as well as its potential cellular role requires further investigation.

It is unclear how long ubiquitin chains need to be in cells to elicit a p97-dependent signalling response. Studies estimate average chain length to be 2-7 ubiquitin long at steady state (Tsuchiya *et al*, 2018), raising the question of how human p97 would engage and process clients if they are only modified with short ubiquitin chains. Hence, another important finding of our work is that FAF2 restores yeast-like efficiency to the human p97-UN complex and confers the same requirement of minimum five K48-linked ubiquitin as the yeast complex(Twomey *et al*, 2019). The simplest explanation for this is that FAF2 works as a ubiquitin binding protein, and indeed an obvious candidate is the UBA domain in FAF2. Surprisingly, we find that the UBA domain is dispensable for its activity. Instead, we identify the activation motif to be a small stretch of ∼25 residues comprising the minimal functional motif. The key residues and conformation of this region were used to develop de novo binders, which successfully mimic the function of FAF2 bypassing the need for p97 association via UBX domain. In line with our observation, deletion of the UBA domain does not impact FAF2’s cellular roles in pexophagy (Koyano *et al*, 2024), lipid droplet biogenesis (Kim *et al*) and stress granule disassembly (Gwon *et al*, 2021). However, the region in FAF2, referred as the activation motif in our study, plays a critical role in these p97 mediated processes.

Intriguingly, we also discover that the region in and around the activation motif of FAF2 is highly unstructured in its apo state and adopts a secondary structure only when bound to UFD1 and p97. This disorder-order transition likely results from multivalent interactions of FAF2 with UFD1 and p97 and this helical conformation is important for ubiquitin engagement by p97-UN-FAF2 complex. Hence, it is unlikely that this unstructured region of FAF2 on its own functions as a UBD, as suggested by a recent study (Huo *et al*, 2025), since the residues important for interaction are unlikely to be correctly positioned. The requirement for the activation motif to be helical is further strengthened by the success of the designed de novo activators where an α-helical structure was enforced during motif scaffolding, resulting in a more stable helical conformation around this region. Furthermore, Huo et al suggest FAF2 (275-349) harbouring the activation motif to function as a UBD by binding polyUb chains of 3 or more ubiquitin. However, we do not observe any binding between FAF2 and polyUb in absence of the UBA domain. Furthermore, the predicted FAF2–Ub3 binding model is sterically incompatible with the p97-UN-FAF2 complex, as the UFD1 UT3 domain occupies the same site on FAF2 as the predicted ubiquitin-binding site. Hence, our data rule out a model in which FAF2-mediated activation of p97-UN complex is driven via a standalone UBD in this region.

Given our single turnover unfolding assay only measures, engagement and processing of ubiquitylated substrates but not release of unfolded peptide from the p97 complex, we think substrate engagement would be a rate-limiting step in our experimental set-up. Importantly, we believe that both FAF2 and the engineered activators use a common mechanism stabilizing interactions between UFD1 and the Ub proximal to the initiator ubiquitin, thereby overcoming this rate-limiting step. This would promote stable engagement of the initiator ubiquitin within the NPL4 groove, facilitating the commitment step prior to ATPase -driven threading and unfolding. Notably, human NPL4 undergoes dynamic seesaw-like conformational changes, with only a specific conformation that aligns NPL4 groove on top of the D1 pore to allow substrate entry, a feature that is unique to the human complex (Pan *et al*, 2021b). Whether FAF2 stabilizes this productive conformation through its interactions with UFD1 and proximal ubiquitin remains an important question for future investigation. However, FAF2 requires its UBX domain for this function. In contrast, our proof-of-principle de novo activators that mimic the FAF2 activation motif to bridge UFD1 UT3 and ubiquitin interactions lack p97 anchoring via the UBX domain. It is likely that additional activator interactions with substrate-engaged p97-UN or stronger affinity for UFD1 compensate for lack of p97 anchoring via the UBX domain.

Activation of p97 would be a desirable strategy to disassemble misfolded protein aggregates that are cytotoxic and a common feature of several proteinopathies. Indeed, p97 can disassemble protein aggregates both in vitro and in cells and loss of p97 activity is associated with vacuolar tauopathy and ALS (Darwich *et al*, 2020; Meyer & Weihl, 2014). The p97 activators reported to date have emerged from screens and increase ATPase activity allosterically by targeting regulatory sites outside the D2 domain or through unknown mechanisms (Jones *et al*, 2024; Phan *et al*; Figuerola-Conchas *et al*, 2020; Wrobel *et al*, 2022). Nevertheless, these activators enhance clearance of disease-causing aggregates from cell models. Our rational design of p97 activators that only work in a UFD1-dependent manner, could provide advantages over non-specific activators as they only enhance unfolding of p97-UN complexes. While being a proof-of-principle demonstration, further optimization of these de novo activators and development of peptides and small molecule activators mimicking FAF2-mediated activation represents an attractive opportunity to modulate cellular proteostasis.

## Methods

### Plasmids

A list of plasmids used in this study can be found in **Table 1**. All plasmids are available upon request by contacting Y.K. or the Medical Research Council Protein Phosphorylation and Ubiquitylation Unit (MRC PPU) Reagents and Services. at https://mrcppureagents.dundee.ac.uk/.

**Table 1.** Plasmids used in this study.

| Recombinant DNA | Reference or source | Identifier or catalogue number |
| --- | --- | --- |
| pET-28a-6His-T7-UFD1_Binder01 | This study (*synthesized by Twist Bioscience*) | DU83270 |
| pET-28a-6His-T7-UFD1_Binder04 | This study (*synthesized by Twist Bioscience*) | DU83273 |
| pET-28a-6His-T7-UFD1_Binder05 | This study (*synthesized by Twist Bioscience*) | DU83274 |
| pET-28a-6His-T7-UFD1_Binder06 | This study (*synthesized by Twist Bioscience*) | DU83275 |
| pET-28a-6His-T7-UFD1_Binder07 | This study (*synthesized by Twist Bioscience*) | DU83276 |
| pET-28a-6His-T7-UFD1_Binder10 | This study (*synthesized by Twist Bioscience*) | DU83279 |
| pET-28a-6His-T7-UFD1_Binder12 | This study (*synthesized by Twist Bioscience*) | DU83281 |
| pET-28a-6His-T7-UFD1_Binder13 | This study (*synthesized by Twist Bioscience*) | DU83282 |
| pET-28a-6His-T7-UFD1_Binder14 | This study (*synthesized by Twist Bioscience*) | DU83283 |
| pET-28a-6His-T7-UFD1_Binder15 | This study (*synthesized by Twist Bioscience*) | DU83284 |
| pET-28a-6His-T7-UFD1_Binder16 | This study (*synthesized by Twist Bioscience*) | DU83285 |
| pET-28a-6His-T7-UFD1_Binder17 | This study (*synthesized by Twist Bioscience*) | DU83286 |
| pET-28a-6His-T7-UFD1_Binder18 | This study (*synthesized by Twist Bioscience*) | DU83287 |
| pET-28a-6His-T7-UFD1_Binder19 | This study (*synthesized by Twist Bioscience*) | DU83288 |
| pET-28a-6His-T7-UFD1_Binder20 | This study (*synthesized by Twist Bioscience*) | DU83289 |
| pFBHTB-6His-UBE1 | MRC-PPU Reagents | DU3026 |
| pET28a(+)-6His-UBE2R1 | MRC-PPU Reagents | DU4317 |
| pET24-Ub | MRC-PPU Reagents | DU20027 |
| pET24-Ub<sup>K48R K63R</sup> | MRC-PPU Reagents | DU23835 |
| pET24-Ub—<sup>1</sup>Ub<sup>1-72, K48R K63R</sup> | MRC-PPU Reagents | DU67512 |
| pET28a-GP78-UBE2G2 | (Blythe *et al*, 2017) | N/A |
| pGEX6P1-GST-3C-UCHL5 | MRC-PPU Reagents | DU12810 |
| pET156P-6His-UBCH5c | MRC-PPU Reagents | DU15703 |
| pET28a-6-His-Ub-mEOS3.2 | This study |  |
| pFL-6His-p97 | This study |  |
| pK27SUMO-14His-Smt3-UFD1 | MRC-PPU Reagents | DU73315 |
| pET28c-NPL4 | MRC-PPU Reagents | DU73317 |
| pK27SUMO-14His-Smt3-Cdc48 | This study |  |
| pK27SUMO-14His-Smt3-Ufd1 | This study |  |
| pK27SUMO-14His-Smt3-Npl4 | This study |  |
| pGEX 3C Halo TEV NSFL1C | MRC-PPU Reagents | DU76358 |
| pK27SUMO-14His-Smt3-FAF1 | MRC-PPU Reagents | DU73318 |
| pK27SUMO-14His-Smt3-FAF2<br>Δ90-118 | MRC-PPU Reagents | DU73323 |
| pK27SUMO-14His-Smt3-UBXN7 | MRC-PPU Reagents | DU73329 |
| pK27SUMO-14His-Smt3-UBXN1 | MRC-PPU Reagents | DU73328 |
| pK27SUMO-14His-Smt3-FAF2<br>Δ137 | MRC-PPU Reagents | DU73324 |
| pK27SUMO-14His-Smt3-FAF2<br>Δ90-118, Δ297-349 | MRC-PPU Reagents | DU73326 |
| pK27SUMO-14His-Smt3-FAF2<br>Δ90-118, Δ 358-441 | MRC-PPU Reagents | DU73327 |
| pK27SUMO-14His Smt3-FAF2<br>Δ90-118-3C-Twinstrep | MRC-PPU Reagents | DU76587 |
| pK27SUMO-14His Smt3-FAF2<br>Δ137-3C-Twinstrep | MRC-PPU Reagents | DU76588 |
| pK27SUMO-14His-Smt3-FAF2<br>Δ90-118 R290E | This Study |  |
| pK27SUMO-14His-Smt3-FAF2<br>Δ90-118 R283E | This Study |  |
| pK27SUMO-14His-Smt3-FAF2<br>Δ90-118 Q287 | This Study |  |
| pK27SUMO-14His-Smt3-FAF2<br>Δ90-118 D304A | This Study |  |
| pK27SUMO-14His-Smt3-FAF2<br>Δ90-118 D294E | This Study |  |
| pK27SUMO-14His-Smt3-FAF2<br>Δ90-118 E307K | This Study |  |
| pK27SUMO-14His-Smt3-FAF2<br>Δ90-118 K308E | This Study |  |
| pK27SUMO-14His-Smt3-FAF2<br>Δ90-118 Y297A | This Study |  |
| pK27SUMO-14His-Smt3-FLAG-UFD1 | This Study |  |
| pK27SUMO-14His-Smt3-FLAG-UFD1 Y162A | This Study |  |
| pK27SUMO-14His-Smt3-FLAG-UFD1 R147E | This Study |  |
| pK27SUMO-14His-Smt3-FLAG<br>UFD1 L53A | This Study |  |

### Protein expression

For recombinant protein expression in *E. coli*, each plasmid was transformed into *E. coli* Rosetta (DE3) cells, which were grown in LB medium supplemented with 50 µg ml⁻¹ kanamycin or 100 µg ml⁻¹ ampicillin, as applicable. Protein expression was induced at an OD_600_ of ∼0.8 with 0.5 mM IPTG, followed by overnight incubation at 18 °C. For expression of WT, mutant ubiquitin and mutant M1-Ub_2_ chains autoinduction medium was used instead of LB, and cultures were incubated at 25 °C for 24 h. Cells were harvested by centrifugation at 4,000 rpm for 20 min at 4 °C.

Human p97 was expressed in insect (Sf9) cells using the baculovirus expression system. Sf9 cells were infected with baculovirus encoding 6xHis-p97, and protein expression was carried out at 27 °C for approximately 72 h, until YFP signal indicated >60% infection. Cells were harvested by centrifugation at 2,500 rpm for 30 min at 4 °C.

### Purification of human p97-UFD1-NPL4 unfoldase complex

For purification of 6×His-tagged human p97 from Sf9 cells, cell pellets were resuspended in ice-cold lysis buffer A (50 mM HEPES pH 7.5, 200 mM NaCl, 10 mM MgCl₂, 5% glycerol, 1 mM ATP, 0.5 mM TCEP) supplemented with cOmplete EDTA-free protease inhibitor cocktail (Roche) and DNase (Sigma). Cells were lysed using a Dounce homogenizer with 20–30 strokes, followed by centrifugation at 20,000 rpm for 30 min. The clarified lysate was incubated with Ni–NTA agarose beads (Amintra) for 1h at 4 °C, and bound protein was eluted with lysis buffer supplemented with 350 mM imidazole. The protein was further purified using a ResourceQ anion-exchange column (6 ml), eluting with a linear NaCl gradient from 150 to 750 mM. Elution fractions were concentrated, and the p97 hexamer was isolated by size-exclusion chromatography on a Superose 6 Increase 3.2/300 column in SEC buffer A (60 mM HEPES pH 7.5, 200 mM NaCl, 0.5 mM TCEP, 10 mM MgCl₂, 5% glycerol).

To reconstitute the UFD1–NPL4 complex, cell pellets expressing each component were resuspended in an equal volume of lysis buffer B (50 mM Tris pH 7.5, 300 mM NaCl, 0.5 mM TCEP, 20 mM imidazole) supplemented with protease inhibitor. Lysates containing His–Smt3–tagged UFD1 and untagged NPL4 were mixed at a 1:2 ratio. Cells were lysed by sonication, and the lysate was clarified by centrifugation at 20,000 rpm for 30 min. The supernatant was incubated with Ni–NTA agarose resin, after which the beads were washed and the UFD1–NPL4 complex was eluted using lysis buffer B supplemented with 350 mM imidazole. The eluted protein was dialyzed overnight into imidazole-free buffer in the presence of His-tagged Ulp1 protease to cleave the His–Smt3 tag from UFD1. The cleaved tag and Ulp1 protease were removed by incubation with Ni–NTA agarose, and the resulting flow-through was concentrated and applied to a Superdex 200 16/600 size-exclusion column equilibrated in SEC buffer A. Fractions containing both UFD1 and NPL4 were pooled, concentrated, and flash frozen.

### Purification of additional p97 cofactors

All p97 cofactors were cloned with an N-terminal 14xHis or GST tag for affinity purification. Briefly, cell pellets were lysed with lysis buffer B supplemented with a protease inhibitor cocktail, followed by sonication and centrifugation at 20,000 rpm. The clarified lysate was incubated with either Ni-NTA agarose resin for His tagged proteins or Glutathione Sepharose 4B resin (Amintra) for GST tagged cofactors for 1 hour, and bound proteins were eluted with lysis buffer containing 350 mM imidazole. The His-Smt3 tag was removed by incubation with Ulp-1 protease. The GST tag was cleaved by incubation with GST tagged TEV protease. Following tag cleavage, the cofactors were further purified on a ResourceQ anion-exchange column (6 mL) and eluted using a linear NaCl gradient from 100 mM to 750 mM. Finally, the pooled fractions were subjected to size-exclusion chromatography on a Superdex 75 16/600 column equilibrated in SEC buffer B (60 mM HEPES pH 7.5, 200 mM NaCl, 0.5 mM TCEP) to isolate the purified protein.

### Purification of ubiquitin

For purification of wild-type and mutant ubiquitin, as well as mutant M1-Ub₂ chains, cell pellets were resuspended in 20 ml of Ub lysis buffer (1 mM EDTA, 1 mM AEBSF, and 1 mM benzamidine). Cells were lysed by sonication, and the lysate was clarified by centrifugation at 20,000 rpm for 30 min. The pH of the supernatant was adjusted to 4.5 by adding 7% perchloric acid, followed by overnight incubation. The mixture was clarified again by centrifugation at 20,000 rpm for 30 min at 4 °C. Ubiquitin was purified using cation-exchange chromatography on a Resource S column (6 ml) in 50 mM sodium acetate pH 4.5, eluting with a NaCl gradient. The pH of the elution fractions was adjusted to 7.5 by adding 100 mM Tris-HCl pH 8.5, after which the protein was concentrated and finally buffer-exchanged into 50 mM HEPES pH 7.5, 150 mM NaCl.

### Preparation of ubiquitylated EOS

To prepare the ubiquitylated mEOS3.2, His tagged Ub-mEOS3.2 and His-tagged GP78-UBE2G2 fusion were purified from *E.coli* and His-tagged UBE1 was purified from insect cells. For bacterial expression, briefly, cells pellets were lysed in lysis buffer B supplemented with protease inhibtors, followed by sonication and centrifugation to obtain the clarified lysate. The lysate was incubated with Ni-NTA agarose beads for 1 hour and the beads were washed with lysis buffer. The bound protein was eluted with 350 mM imidazole, For GP78-UBE2G2, the His-tag was cleaved by incubating the protein with His-TEV protease for overnight. The cleaved protein and cleaved tag were removed by reverse IMAC (Immobilized Metal Affinity Chromatography). For Ub-Eos, the His tag was retained. The proteins were further purified using to size-exclusion chromatography on a Superdex 75 16/600 column equilibrated with SEC buffer C (50 mM HEPES pH 7.5, 150 mM NaCl, 0.5 mM TCEP).

UBE1 was expressed in Sf9 cells using the baculovirus expression system at 27 °C for approximately 72 h. Cells were harvested by Centrifuge at 2500 rpm for 30 mins. and re-suspend in lysis buffer (50 mM HEPES pH 7.5, 150 mM NaCl, 0.5 mM TCEP, 1mM AEBSF, 1mM Benzamidin). The lysate was passed through a dounce homogenizer 12 times to thoroughly lyse the cells. After clarification at 20,000 rpm for 30 minutes 20 mM imidazole was added, and the lysate was incubated for 1 hour with NiNTA resin. The resin was washed extensively, and bound protein was eluted from the beads using lysis buffer supplemented with 350 mM imidazole. The purified UBE1 was dialysed over-night in presence of His-TEV to cleave the tag and further purified using a Superdex 200 16/600 size-exclusion column equilibrated with in SEC buffer C.

Ubiquitylation of the Ub-Eos was carried out as described before (Blythe *et al*, 2017). 10 μM His-tagged Ub-Eos, 1 μM UBE1, 30 μM UBE2G2-gp78 and 400 μM ubiquitin was incubated in reaction buffer (20 mM Hepes, 150 mM NaCl, 0.1 mM TCEP) at 37°C. Ubiquitin was added slowly for during the first 6 hours of the reaction and the reaction was continued overnight. The ubiquitylated Eos were purified by incubation with Ni-NTA agarose resin equilibrated with reaction buffer supplemented with 10 mM imidazole. The beads were washed and bead bound substrate was eluted with 300 mM imidazole. The purified substrate was loaded onto a Superose 6 Increase 3.2/300 column in SEC buffer (20 mM Hepes, 150 mM NaCl, 0.1 mM TCEP) to separate the substrate based on Ub chain length. Substrates were pooled according to the average chain length and segregated into Ub^BT^, Ub^S^ and Ub^L^ substrates. The substrates were photoconverted using a 405 nM laser until 50% or more photoconversion was achieved.

### Preparation of Ub_6_-Eos substrate

K48 linked tetra Ub chains were synthesized by 1.5 mM Ub, 1 μM UBE1 and 25 μM UBE2R1 in ligation buffer (40 mM Tris-HCl (pH 7.5), 10 mM MgCl₂, and 10 mM ATP) at 37 °C for 4–6 h. The chains were purified using Resource S column (6 ml) equilibrated in 50 mM sodium acetate (pH 4.5) and eluted with a linear NaCl gradient. For C-terminal capping, tetraubiquitin was incubated with 2 μM UBE1, 10 μM UBE2D3, and 100 μM L-lysine in ligation buffer at 37 °C for 16 h and purified by cation-exchange chromatography under identical conditions. N-ter capped diUb was prepared by using Ub ^K48R, K63R^ mutant and M1-Ub_2_ (Ub ^K48R, K63R^, Ub^1-72^. Ligation and purification were carried out as described previously (Lange *et al*, 2024). The capped diUb was conjugated to the protected tetraUb using 2 μM UBE1 and 20 μM UBE2R1 in ligation buffer. After 24 h, 1 μM GST-UCHL5 was added to deprotect the C-terminus, and the reaction was incubated at 30 °C for an additional 2 h. The resulting K48-linked hexamer with capped distal Ub (Ub ^K48R, K63R^) was purified using Resource S column equilibrated in 50 mM sodium acetate (pH 4.5). The capped hexamer was subsequently loaded onto Ub-mEOS using 2 μM UBE1 and 20 μM UBE2R1 and Ub_6_-Eos substrate was purified by cation-exchange chromatography using conditions as described before.

### Synthesis of K48-linked polyUb chains

K48-linked Ub chains were assembled from 1.5 mM Ub in 40 mM Tris-HCl (pH 7.5), 10 mM MgCl₂, and 10 mM ATP at 37 °C for 4–6 h. Formation of K48 linkage was catalyzed by 1 μM UBE1 and 25 μM UBE2R1. For assembly of Ub₆ or Ub₈ chains, diUb was used as the starting material. Ub chains were separated by length using cation-exchange chromatography on a Resource S column (6 ml) equilibrated in 50 mM sodium acetate (pH 4.5) and eluted with a linear NaCl gradient. Fractions were pooled according to chain length, concentrated using 3-kDa or 10-kDa MWCO centrifugal filter units (Amicon), and buffer-exchanged into 50 mM HEPES (pH 7.5), 150 mM NaCl.

### Single turnover unfolding assay

Single turnover unfolding assays were carried out by incubating 20 nM photoconverted red mEOS3.2 substrate modified with Ub chains of various lengths, 400nM p97, 400 nM UFD1-NPL4 and 1.6 μM of additional cofactors (p47, UBXN2B, UBXN7, FAF1, FAF2 and UBXN1) or 1.6 μM of *de novo* binders in assay buffer (20 mM HEPES pH 7.5, 150 mM KCl, 10 mM MgCl_2_, 1 mM TCEP,1 mg/ml BSA) at 37°C for 10 mins. The unfolding reaction was initiated by addition of 5 mM ATP. Fluorescence was monitored using BMG Labtech CLARIOstar plate reader at 37°C. Unfolding was measured by monitoring loss of fluorescence at 590nM upon excitation at 540 nM. The percentage of folded substrate remaining at each time point was calculated by normalizing to the fluorescence at time 0 and to changes in fluorescence observed for substrate-only control. The unfolding rate (*k*, sec^-1^) was calculated by fitting the unfolding traces to single exponential decay. The initial unfolding rate was determined by linear regression of the fluorescence decay data from the first 2 minutes of the reaction. Rates are reported as the slope of the linear fit in units of % fluorescence decay per second.

EC_50_ values for cofactor-mediated activation were determined by performing single-turnover unfolding assays with varying concentrations of FAF1, FAF2, or *de novo* binders while maintaining constant concentrations of p97, UFD1–NPL4 and ubiquitylated fluorescence substrate as described above. Initial (∼ first 2 min) rates of unfolding were plotted as a function of log_10_ or log_2_ cofactor concentration and fit to a four-parameter logistic curve in GraphPad Prism to extract EC_50_ values.

### ATPase assay

ATPase assay was carried out using 10 nM p97, 100 nM UFD1-NPL4, 100 nM FAF2 and 100 nM Ub^L^ substrate or 500 nM free K48-linked Ub chains of specific length in assay buffer (20 mM HEPES pH 7.5, 150 mM KCl, 10 mM MgCl_2_, 1 mM TCEP, 0.1 mg/ml BSA). Proteins were mixed and incubated in 384 well microplate (Grenier 781101) at 37°C for 10 minutes followed by addition of 200 μM ATP to initiate the reaction. Reactions were incubated at 37°C for 30 minutes to allow ATP hydrolysis. The reactions were stopped by addition of 50 µl of BIOMOL Green (Enzo Life Sciences) and further developed at room temperature for 1 hour. Absorbance was recorded at 620 nM using BMG Labtech CLARIOstar plate reader. Inorganic phosphate released from each reaction was quantified based on a phosphate standard curve. Relative ATPase activity was determined by normalizing the measured activity of each sample to that of the control.

### In-vitro pulldown of p97-UN-FAF2 complex

Recombinant FAF2-Twin Strep tagged, UFD1-NPL4 and p97 (hexamer) proteins were preincubated on ice for 30 min in binding buffer (20 mM HEPES pH7.5, 150 mM NaCl, 10 mM MgCl_2_, 1 mM ATP and 0.5 mM TCEP) at a final concentration of 5 µM in a 1:1 molar ratio. The resulting complexes were incubated with 10 µL of pre-equilibrated Pierce™ High-Capacity Streptavidin Agarose (Thermo Fischer) in pulldown buffer (20 mM HEPES pH 7.5, 150 mM NaCl, 10 mM MgCl_2_, 0.1 mM ATP, 0.5 mM TCEP and 0.5% NP-40) on a roller at 4°C for 1 hour. Beads were washed six times with ice-cold pulldown buffer. Bead bound protein was eluted with 2× LDS sample buffer and analyzed by SDS–PAGE followed by oriole stain.

For FLAG pulldown, recombinant FLAG-tagged UFD1-NPL4 complex, FAF2 and p97 were incubated under the same conditions as described above. The resulting complexes were incubated with 20 µL of pre-equilibrated anti-FLAG M2 Affinity Gel (Sigma) in pulldown buffer for 1 hour. Beads were washed six times, and the bound complex was eluted with 3x Flag peptide prepared in pulldown buffer. The input and pulldown samples were analyzed by SDS–PAGE followed by Oriole stain.

### In-vitro pulldown of ubiquitin

Twin strep-tagged FAF2 (WT and ΔUBA), K48-linked Ub trimer (Ub3), and hexamer (Ub6) chains were mixed in binding buffer (20 mM HEPES pH7.5, 150 mM NaCl, and 0.5 mM TCEP) at a final concentration of 2.5 µM in a 1:1 molar ratio and were incubated with 10 µL of pre-equilibrated Pierce™ High Capacity Streptavidin Agarose (Thermo Fischer) in pulldown buffer (20 mM HEPES pH 7.5, 150 mM NaCl, 0.5 mM TCEP and 0.3% DDM) on a roller at 4°C for 1 hour. Beads were washed three times with ice-cold pulldown buffer. Bead-bound protein was eluted with 2× LDS sample buffer and analyzed by SDS–PAGE followed by oriole stain.

### HDX-MS sample preparation

5 µM p97/UFD1-NPL4 with 1 mM ATPγS was incubated with or without 5 µM FAF2, or for FAF2 Apo Conditions 5 µM FAF2 alone, for 30 minutes in Protein Dilution Buffer (20 mM Tris pH 7.5, 150 mM NaCl, 10 mM MgCl_2_, 0.5 mM TCEP) on ice. 5 µl of this sample was diluted with 45 µL of Deuteration Buffer (20 mM Tris pH 7.5, 150 mM NaCl, 10 mM MgCl_2_, 0.5 mM TCEP, 94.6% D_2_O) (final D_2_O concentration = 85.1%) for timepoints 3/30/300/3000 s, before being quenched with 20 µl of ice-cold Quench Solution (6 M Urea, 2% Formic Acid), and being snap frozen in liquid nitrogen and stored at –70°C. A further time point, 0.3 s, was achieved by incubating both the sample and deuteration buffer on ice and then conducting a 3 s exchange reaction. Each exchange reaction was conducted independently four times.

### HDX-MS data acquisition

Samples were rapidly thawed at room temperature and then injected into automated HDX-MS fluidics and UPLC manager system (Waters). Samples were loaded into a 50 µl loop and then subsequently digested using a Waters Enzymate BEH Pepsin Column (Part No. 186007233) in a 0.1% Formic Acid solution with a flow rate of 200 µl/min at 20°C. Peptic peptides then flowed onto a Waters ACQUITY UPLC BEH C18 VanGuard Pre-Column at 1°C (Part No. 186003975). After digestion, the flow path was changed to elute the peptides via a Waters ACQUITY UPLC BEH C18 1.7 µm 1.0 × 100 mm reverse phase column (Part No. 186002346). A gradient from 0-85% 0.1% Formic Acid/ Acetonitrile, conducted at 1°C with a 40 µl/min flowrate, was used to elute deuterated peptides which were then ionised using an ESI source. Mass spectrometry data were collected using a Waters Select Series cIMS instrument, from a 50–2,000 m/z range with the instrument in HDMSe mode. A single pass of the cyclic ion mobility separator (with a cycle time of 47 ms) was conducted. A blank sample of protein dilution buffer with quench was run between samples, and carryover was routinely checked to be <1% intensity of the prior sample.

### HDX-MS data analysis

Peptide sequence identification was conducted using Protein Lynx Global Server (Waters), searching against a database of the protein complex, common contaminants, porcine pepsin, and previous project’s sequences. Search criteria include S/T/Y phosphorylation. Minimum inclusion criteria were a minimum intensity of 5000 counts, minimum sequence length 5, maximum sequence length 35, a minimum of three fragment ions, a minimum of 0.1 products per amino acid, a minimum score of 5.0, a maximum MH+Error of 10 ppm. Subsequent analysis and determination of deuteration values conducted using HDExaminer (Sierra Analytics/Trajan). Experimental design, data acquisition, analysis and reporting are in line with the community agreed recommendations (Masson *et al*, 2019).

### Cross-linking mass spectrometry (XL-MS) sample preparation

Cross-linking reactions were optimised by varying disuccinimidyl sulfoxide (DSSO, Thermo Fisher Scientific) (Kao *et al*, 2011) concentration, protein concentration, and cross-linking time, and analysed by SDS-PAGE and mass photometry. 5 μM p97 hexamer was incubated for 30 minutes at 25°C in 50 mM HEPES pH 7.5, 5 mM MgCl_2_, 1 mM ATP𝛾S containing (as appropriate) 10 μM UFD1-NPL4, 10 μM FAF2, and 10 μM K48-linked Ub_6-8_ ubiquitin chain. Freshly prepared DSSO was added to a final concentration of 600 μM and the cross-linking reaction allowed to proceed for a further 15 min at 25°C before quenching with 80 mM Tris-HCl pH 8.8. Four independent replicates were performed for each condition.

Cross-linked samples were denatured and reduced in 1% (w/v) SDS, 10 mM DTT for 5 min at 95°C. Ammonium bicarbonate buffer (pH 8) was added to a final concentration of 10 mM and proteins were alkylated with 50 mM iodoacetamide at room temperature for 30 min in the dark. Trichloroacetic acid (TCA) was added to a final concentration of 20% (v/v) to precipitate protein, and samples were incubated on ice for 15 min before centrifugation at 17,000 x *g* for 10 min at 4°C. Proteins were resuspended in 10% (v/v) TCA and pelleted once more by centrifugation at 17,000 x *g* for 5 min. Pellets were washed 3 times with MS-grade water (17,000 x *g* for 5 min at 4°C) and resuspended in 30 μL of a denaturing solution containing 8 M urea and 20 mM DTT. Following incubation for 10 min at 30°C, each replicate was further split into 3 x 10 μL aliquots and 90 μL of 25 mM ammonium bicarbonate buffer added to adjust the urea concentration to 0.8 M. All samples were digested for 20 hours with 2 ng/μL MS-grade trypsin (Thermo Fisher Scientific) at 30°C. Additionally, 4 hours after the beginning of this incubation, double digests were created by adding to the auxiliary aliquots created for each initial replicate either Asp-N endoprotease (Promega) to a final concentration of 0.2 ng/μL or Glu-C endoprotease (Promega) to a final concentration of 1 ng/μL (thereby creating 3 digestion conditions in order to maximise sequence coverage).

### LC-MS/MS analysis of crosslinked peptides

Liquid chromatography tandem mass spectrometry (LC-MS/MS) experiments were performed on an UltiMate 3000 RSLC nano-HPLC system coupled to an Orbitrap Lumos mass spectrometer. 5 µl solution from each digested cross-linked sample was loaded onto the nano-HPLC system individually. Peptides were trapped by a precolumn (Acclaim PepMapTM 100, C18, 100 µm × 2 cm, 5 µm, 100 A) using aqueous solution containing 0.1% (v/v) TFA. Peptides were then separated by an analytical column (PepMapTM RSLC C18, 75 µm × 50 cm, 2 µm, 100 A) at 45°C using a linear gradient of 3 to 35% solvent B (solution containing 80% ACN and 0.1% FA) over 90 mins, 35 to 85% solvent B over 5 mins, 85% solvent B for 10 min, 85% to 3% solvent B over 1 min and 3% solvent B for a father 8 min. The flow rate was set at 300 nl/min for all experiments. Data were acquired with data-dependent MS/MS mode. For each MS scan, the scan range was set between 375 and 1,500 m/z with the resolution at 120,000 and 300% automatic gain control (AGC) was used. The maximum injection time for each MS scan was 100 ms. Peptides with charge state between 2 and 8 as well as intensity threshold higher than 1.0e+4 were then isolated with a 1.2 Da isolation window sequentially. Stepped HCD with normalized collision energy of 27, 30 and 33% was applied to fragment the isolated peptides. For each MS/MS scan, the resolution was set at 15,000 with a normalized AGC at 200%. The maximum injection time was set at 250 ms. Dynamic exclusion with 60-s duration and 10 ppm window was enabled for the experiment.

### XL-MS data analysis

The .RAW files obtained from LC-MS/MS were converted into .mgf files using the MSConvert tool within the ProteoWizard software suite (Holman *et al*, 2014). The resulting .mgf files were analysed using MeroX to assign and discriminate the cross-linked products (Götze *et al*, 2015). Settings corresponding to peptide lengths ranging from 3 to 50 amino acids containing a maximum of 3 missed cleavages were applied, with the appropriate digestive enzyme(s) considered. Carbamidomethylation at cysteine residues was set as a static modification and oxidation at methione and deamidation at asparagine were included as variable modifications. DSSO was selected as the cross-linker, with its principle specificity set for lysine residues, but also considering reactivity for serine, threonine, tyrosine, and the N-terminus of target proteins (Iacobucci *et al*, 2019; Smith *et al*, 2018). 10 ppm and 20 ppm were used to filter the mass error in precursor ion (MS1) and fragment ion (MS2) scans. Only ions with a signal-to-noise ratio higher than 2 went forward for database searching. RISEUP searching mode was applied, with a minimum of 2 fragments per peptide and a false discovery rate (FDR) cut-off of 5% set for cross-linked peptide identification. The common Repository of Adventitious Proteins (cRAP) was included to increase the size of the decoy database and improve FDR estimation. Data from the three different protease digestion conditions (trypsin alone, trypsin + Asp-N, trypsin + Glu-C) were recombined into a single data set and potential cross-linked peptides with a MeroX score higher than 50 were manually inspected to ensure unambiguous cross-link identification (Iacobucci *et al*, 2018). Filtered cross-linked data was exported from MeroX and uploaded to xiVIEW for visualisation and figure preparation (Combe *et al*, 2024).

### Mass photometry

Proteins were cross-linked with DSSO as described for cross-linking mass spectrometry. Samples were then diluted 50-fold in phosphate buffered saline (PBS) to a final concentration of approximately 100 nM p97 hexamer. Mass photometry was performed using a Refeyn One^MP^ mass photometer (Refeyn Ltd, Oxford, UK) combined with AcquireMP and DiscoverMP software for data acquisition and analysis, respectively (Wu & Piszczek, 2021; Young *et al*, 2018). Microscope coverslips (VWR) and Grace Bio-Labs reusable Culturewell gaskets (Sigma) were cleaned by washing sequentially 3 times with isopropanol and deionised water and then dried with a stream of compressed air. A calibration curve was established consisting of sweet potato β-amylase monomer (56 kDa) and dimer (112 kDa), bovine thyroglobulin dimer (670 kDa), and the lactate dehydrogenase (146 kDa) and apoferritin band 2 (480 kDa) components of Thermo’s NativeMark unstained protein standard. To find focus, 5 µL of PBS was pipetted into a fresh well of the gasket/coverslip assembly and the focal position was identified. 5 µL of cross-linked protein sample was added to the same well to achieve a final working concentration of approximately 50 nM p97 hexamer complex. A 60 s movie was recorded, and the distribution peaks were fitted with Gaussian functions to obtain the average molecular mass of each distribution component.

### Design of UFD1-binding FAF2 helix scaffolds with RFdiffusion

A motif scaffolding approach was taken to design potential p97-activating proteins scaffolded upon the predicted UFD1-binding helix of FAF2. AlphaFold3 (Abramson *et al*, 2024) was used to design a model of the complex between p97 (N-terminal and D1 ATPase domains), NPL4, UFD1, FAF2, and 10 ubiquitin. This model was used as the input for binder generation using RFdiffusion (Watson *et al*, 2023), integrated with ProteinMPNN amino acid prediction (Dauparas *et al*, 2022) and AlphaFold2-based folding and validation (Jumper *et al*, 2021), and run online as a single pipeline via a Google Colab notebook. Residues T286, L289, R290, Q292, Q293, D294, A296, Y297, S300, L301, D304, Q305, K307, E308, and K311 were fixed and scaffolded and various N- and C-terminal extensions of between 5 and 160 amino acids were sampled, with RFdiffusion also inpainting the intervening residues within the FAF2 helical scaffold to implicitly reason over their sequence and better pack against them (Wang *et al*, 2022). This motif scaffolding of the FAF2 helix was performed in the presence of the UT3 domain of UFD1 (amino acids 17-196) and two ubiquitin molecules. While we did experiment with the selection of hotspot residues, none of the high-ranking nor successful designs derived from design trajectories involving hotspots. RFdiffusion settings were left largely as default or Baker Lab recommended settings (50 iterations, 8 ProteinMPNN sequences per design, MPNN sampling temp of 0.1, cysteines excluded from consideration during MPNN step, AF2 + initial guess used to validate protein structures with 3 recycle steps).

In total 2248 UFD1 binders, representing 281 distinct backbone trajectories, were generated over 16 Colab runtimes. Candidates were filtered based on pAE (<4) and plDDT (>0.85), with pTM scores above 0.75 and RMSD scores less than 2Å also considered desirable. To promote diversity amongst the binder designs only the top ranked ProteinMPNN-generated sequence belonging to each design trajectory was considered. One exception to the filtered designs outlined above was Binder 4 *(Run12_Design8_mpnn5)*, which failed on all metrics but stood out as the only design out of 256 generated during a rather unsuccessful Runtime that was predicted by AlphaFold to bind to UFD1 in the same location as FAF2.

### AlphaFold

AlphaFold3 was used to model p97-UN-FAF2, p97-UN-FAF1 and p97-UN-FAF2-polyUb complex (Abramson *et al*, 2024). From the five predictions generated for each complex, the model with the highest ipTM (interface predicted Template Modeling) score was selected for analysis.

### Bioinformatics analysis

Conservation analysis of UFD1 and FAF2 was performed by Consurf (Ashkenazy *et al*, 2016). Multiple sequence alignments were performed using ClustalW (Sievers *et al*, 2011) and visualized with ESPript3 (Gouet *et al*, 1999). Structural homology search for FAF2 CR was performed using DALI web server (Holm *et al*, 2023).

### Isothermal Titration Calorimetry (ITC)

ITC titration experiments were performed on a MicroCal PEAQ-ITC instrument (Malvern, version 1.29.32) operated with PEAQ-ITC software (Malvern) at 25°C. Before measurement, all proteins were dialysed in ITC buffer (50 mM Tris (pH 7.5), 150 mM NaCl, 0.25 mM TCEP). UFD1 was added to the syringe and titrated into the cell which contained K48-Ub_2_. For titration of UFD1 and K48-Ub_2_, concentration of 199 µM and 30.2 µM was used respectively. Similarly, FAF2 was added to the syringe at a concentration of 141uM and titrated into the cell which contained K48-Ub_2_ at a concentration of 20.1µM. Data were analyzed and titration curves were fitted using MicroCal PEAQ-ITC (Malvern) analysis software and GraphPad Prism 9 was used for figure generation.

## Data analysis and figure preparation

Data were analyzed and plotted using Prism v.10 (GraphPad). Figures were prepared using Adobe Illustrator, and UCSF ChimeraX.

**Supplementary Table S1.**

DataSet 1:
|  | p97 | UFD1 | NPL4 | Overall |
| --- | --- | --- | --- | --- |
| # of Peptides | 323 | 36 | 86 | 445 |
| Coverage (%) | 96.4% | 92.5% | 91.1 | 93.9% |
| Gaps in Coverage | (-6-4),408-453 | 86-97, 225-232, | 336-339,481-494,604-609 |  |
| Av. Length | 15.3 | 17.9 | 17.1 | 15.9 |
| Redundancy | 6.1 | 2.1 | 2.4 | 4.1 |
| Time Points | 0.3s<sup>1</sup> / 3s/ 30s/ 300s/ 3000s |  |  |  |
| Replicates | 3 ND, 3D |  |  |  |
| Replicability<sup>2</sup> | 0.09 Da |  |  |  |
| Significance Threshold<sup>3</sup> | >5%/ >0.5 Da/ passes T-test |  |  |  |

DataSet 2:
|  | FAF2 Δ90-118 |
| --- | --- |
| # of Peptides | 122 |
| Coverage (%) | 87.7% |
| Gaps in Coverage | 81-90, (90-118), 119-140, 300-324 |
| Av. Length | 21.0 |
| Redundancy | 5.8 |
| Time Points | 0.3s<sup>1</sup> / 3s/ 30s/ 300s/ 3000s |
| Replicates | 3 ND, 3D |
| Replicability<sup>2</sup> | 0.2 Da |
| Significance Threshold<sup>3</sup> | >5%/ >1.0 Da/ passes T-test |

1. 0.3 s timepoint created by pipetting 3 s by hand with ice-cold solutions.
2. Mean standard deviation between replicates
3. A peptide must meet all these criteria to have a significant change in solvent exchange. The threshold is set as 5σ.

## Data availability

HDX-MS data has been deposited to the ProteomeXchange Consortium via the PRIDE partner repository with the dataset identifier PXD073479.

## Acknowledgements

We thank members of the Kulathu lab, especially Drs. Rachel O’Dea, Stephen Matthews and Matthew R. McFarland for critical reading of the manuscript and helpful comments. We thank Vincent Brouwer for insightful discussion throughout the project. We acknowledge MRC Reagents and Services, especially Jennifer Thomson for cloning the constructs used in this study. We thank Dr. Linnan Shen for protein reagents and ubiquitin reagents. We thank Dr. Yuko Lam (MRC PPU) for her advice regarding XL-MS and the MRC-PPU mass spectrometry facility (coordinated by Dr. Renata Soares) for technical support. We thank Prof. Martin (UC Berkeley) for providing us the GP78-UBE2G2 construct.

## Author Information

**[Credit](https://credit.niso.org/) role taxonomy**

Conceptualization – PD, YK

Data curation

Funding acquisition – GM, YK

Formal analysis – PD, GM, IK, YK

Investigation – PD, IK, GA, APR, RG

Methodology

Project administration

Resources – AK, RG, GM

Software

Supervision – YK

Validation Visualization – PD, IK

Writing – original draft – PD, YK

Writing – review & editing – PD, IK, GM, YK

## Disclosure and competing interest statement

PD, IK, YK are inventors on a pending patent application filed by the University of Dundee covering the development of p97 activators based on the findings described in this work.

## Funding

This work was funded by UKRI MRC grant MC_UU_00038/3 (YK), ERC Consolidator grant (grant 101002428; YK), Tricia Cohen Memorial Trust PhD studentship (PD). HDX-MS equipment was funded by BBSRC Capital Equipment Fund BB/V019635/1.

## Extended Data

**Figure EV1:**
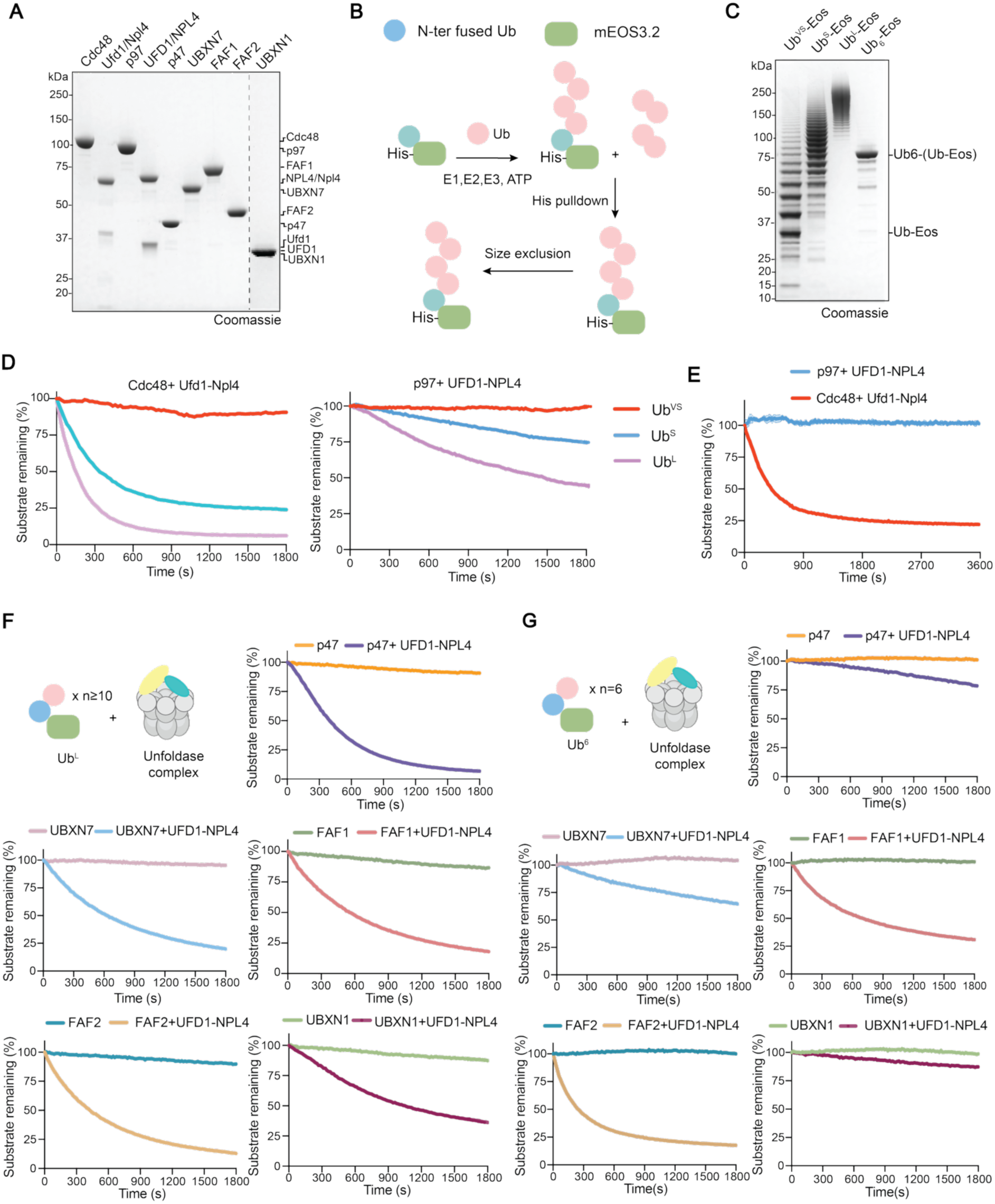
Differential modulation of human p97 activity by UBA–UBX cofactors requires UFD1–NPL4. (A) Coomassie-stained SDS–PAGE showing purified human p97/yeast Cdc48, yeast Ufd1-Npl4 complex, human UFD1–NPL4 complex, and UBA-UBX cofactor proteins used in this study. (B) Schematic representing the overview of the method employed to prepare ubiquitylated reporter Eos substrates. (C) Coomassie-stained SDS–PAGE of K48-linked ubiquitylated Eos substrate of various lengths used in the unfolding assays. (D) Unfolding of Ub^VS^, Ub^S^, and Ub^L^ substrates by yeast Cdc48–Ufd1–Npl4 (left) and human p97–UFD1–NPL4 (right). (E) Comparison of unfolding of Ub_6_-Eos by human p97-UFD1-NPL4 and yeast Cdc48-Ufd1-Npl4 unfoldase complex. (F) Ub^L^ substrate unfolding by human p97 with UBA–UBX cofactors (p47, UBXN7, FAF1, FAF2 or UBXN1), alone or combined with UFD1–NPL4, showing cofactor-specific modulation of activity. (G) Substrate unfolding kinetics of Ub_6_-Eos substrate by human p97 in the presence of UBA-UBX cofactors either alone or in combination with UFD1–NPL4, demonstrating cofactor-specific effects on unfolding depend on ubiquitin chain length on client. For panel (D-G), fluorescence intensity values were normalized to values at time 0 and substrate-only controls. Data shown as mean ± SD (n ≥ 3 technical replicates for D and E, n = 3 independent experiments, each with 3 technical replicates for F and G).

**Figure EV2:**
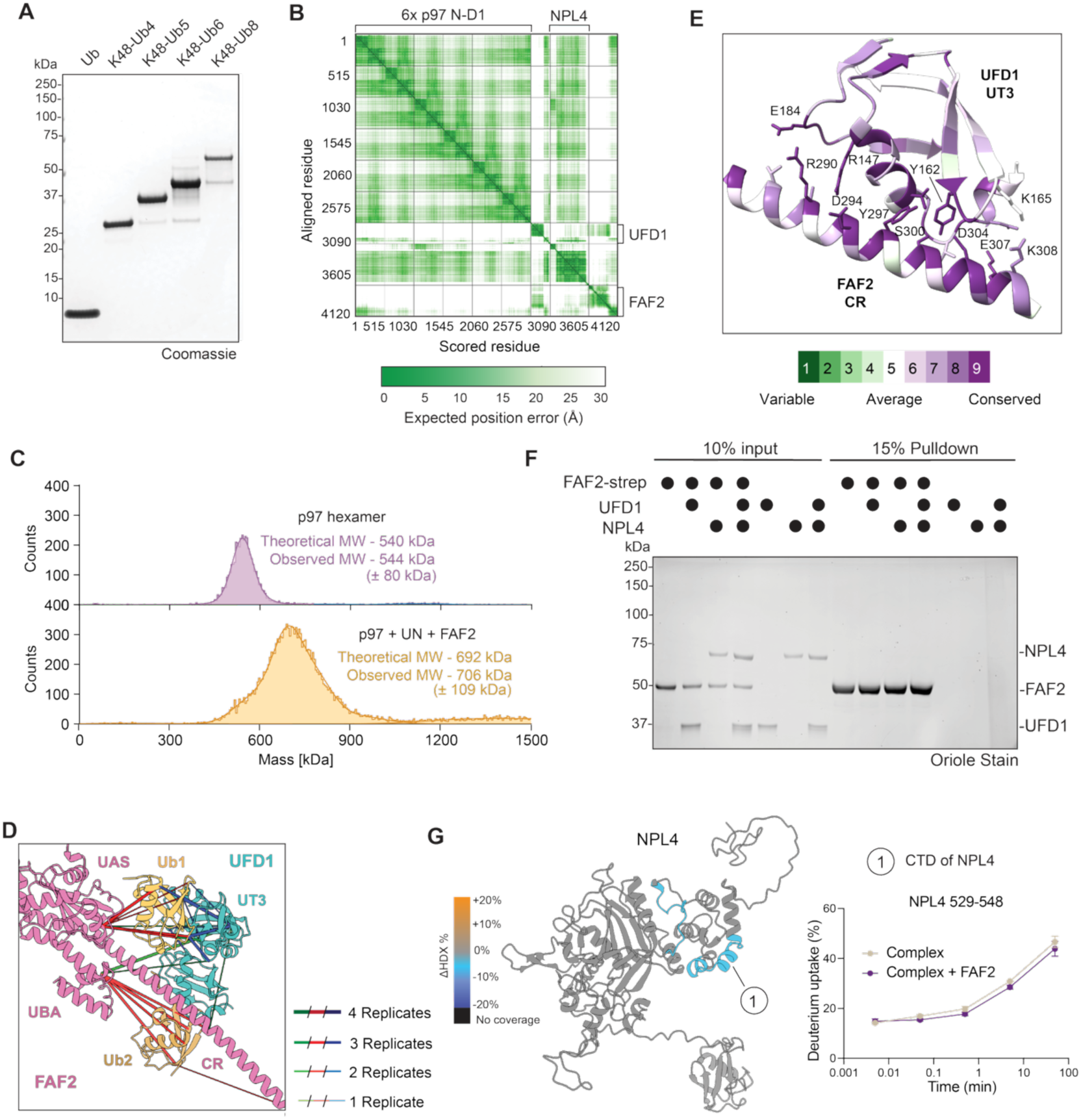
Analysis of p97: UN: FAF2 complex formation. (A) Coomassie-stained SDS–PAGE of K48-linked ubiquitin chains used for substrate-stimulated ATPase activity assay. (B) PAE plot of the represented Alphafold3 model of p97(ND1)-NPL4-UFD1 complex. (C) Mass photometry analysis of crosslinked p97-UFD1-NPL4-FAF2 complex. Conditions include 5 μM p97 hexamer alone (purple), 5 μM p97 hexamer + 10 μM UFD1-NPL4 + 10 μM FAF2 (orange), in the presence of 600 μM DSSO crosslinker. Histograms are representative of n=3 independent experiments. MW, molecular weight. (D) Experimental crosslinks mapped onto the AlphaFold3 model of the p97–UFD1– NPL4–FAF2–polyUb complex. Crosslinked residue pairs are shown as connecting lines (colouring is the same as Figure 2F), supporting the predicted complex architecture. (E) Conservation analysis of FAF2 and UFD1 generated by Consurf from the sequences of 150 eukaryotic species. Conservation of each residue is represented by a score between 1-9, with 1 being the least conserved and 9 being the most conserved. Consurf analysis is then mapped onto the AF3 prediction of p97-UFD1-FAF2 complex and UFD1-FAF2 interface is highlighted. (F) Strep pull-down of C-ter Twin-Strep-tagged FAF2 was performed with UFD1, NPL4, or UFD1–NPL4 complex, followed by Oriole staining. (G) ΔHDX mapping of NPL4 bound to p97 and UFD1 upon FAF2 binding (left). Regions with reduced solvent exchange are shown in blue, increased exchange in orange, no coverage in black, and no significant change (−5% to +5%) in grey. Deuterium uptake kinetics of peptide in the NPL4 CTD domain showing FAF2-dependent protection (right). Data represent mean ± SD (n = 3 independent replicates).

**Figure EV3:**
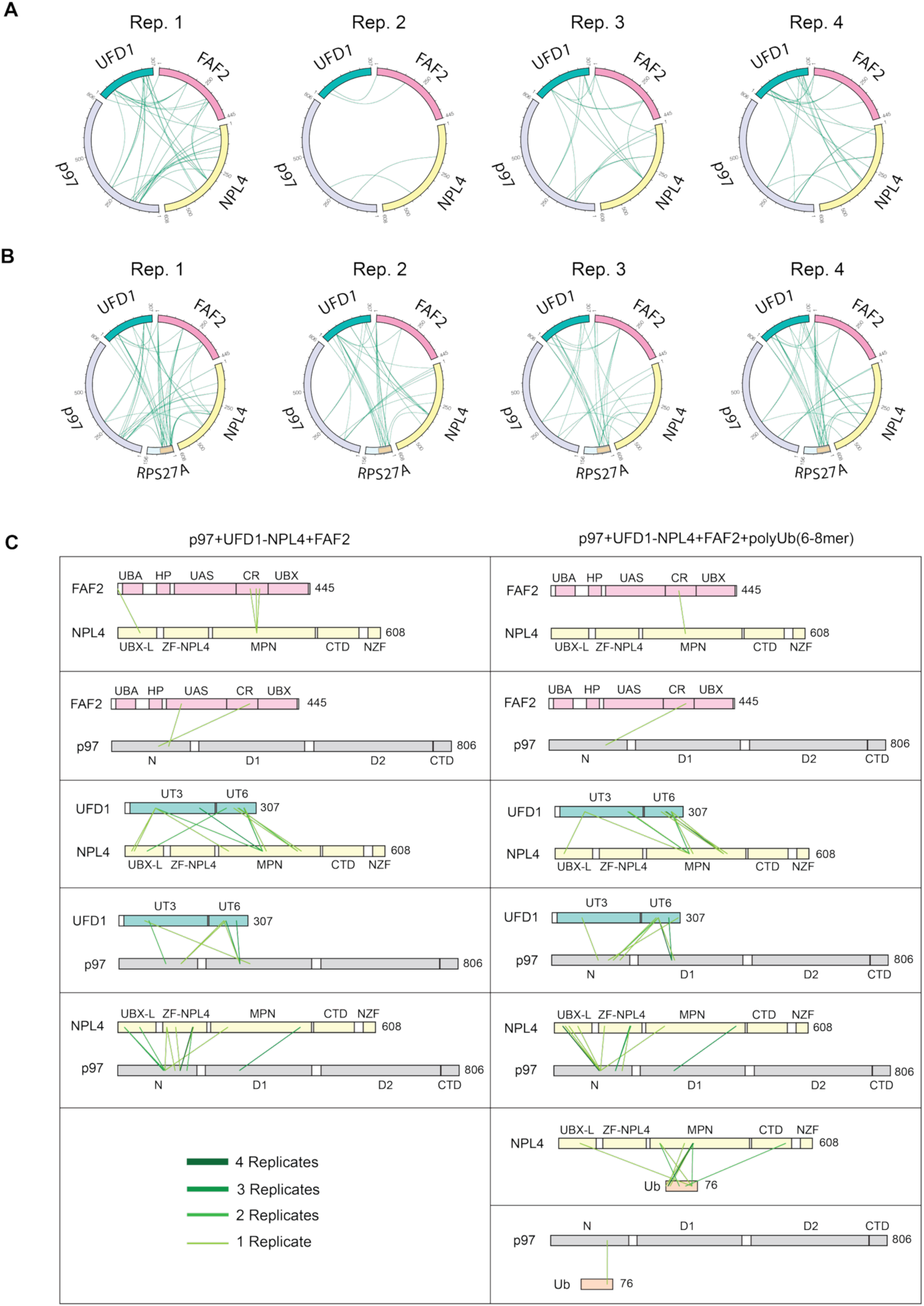
DSSO-mediated cross-linking of p97:UFD1-NPL4:FAF2 complex -/+ ubiquitin. (A) Circular diagrams of DSSO-mediated intermolecular cross-links observed between components of the p97:UFD1-NPL4:FAF2 ternary complex are shown for 4 independent replicates. (B) Circular diagrams of interprotein cross-links observed between components of the quaternary complex formed between p97:UFD1-NPL4:FAF2 and a chain of K48-linked polyubiquitin (denoted as the first 76 amino acids of Ribosomal Protein S27a). (C) The domain architectures of the indicated complex components are shown, highlighting the sites of chemical cross-linking in n = 4 independent replicates. Cross-links are represented by straight lines, the colour and width of which indicates reproducibility across different replicates. Panels on the left indicate the cross-links observed in the absence of ubiquitin, those on the right highlight those cross-links recorded in the presence of poly-Ub.

**Figure EV4:**
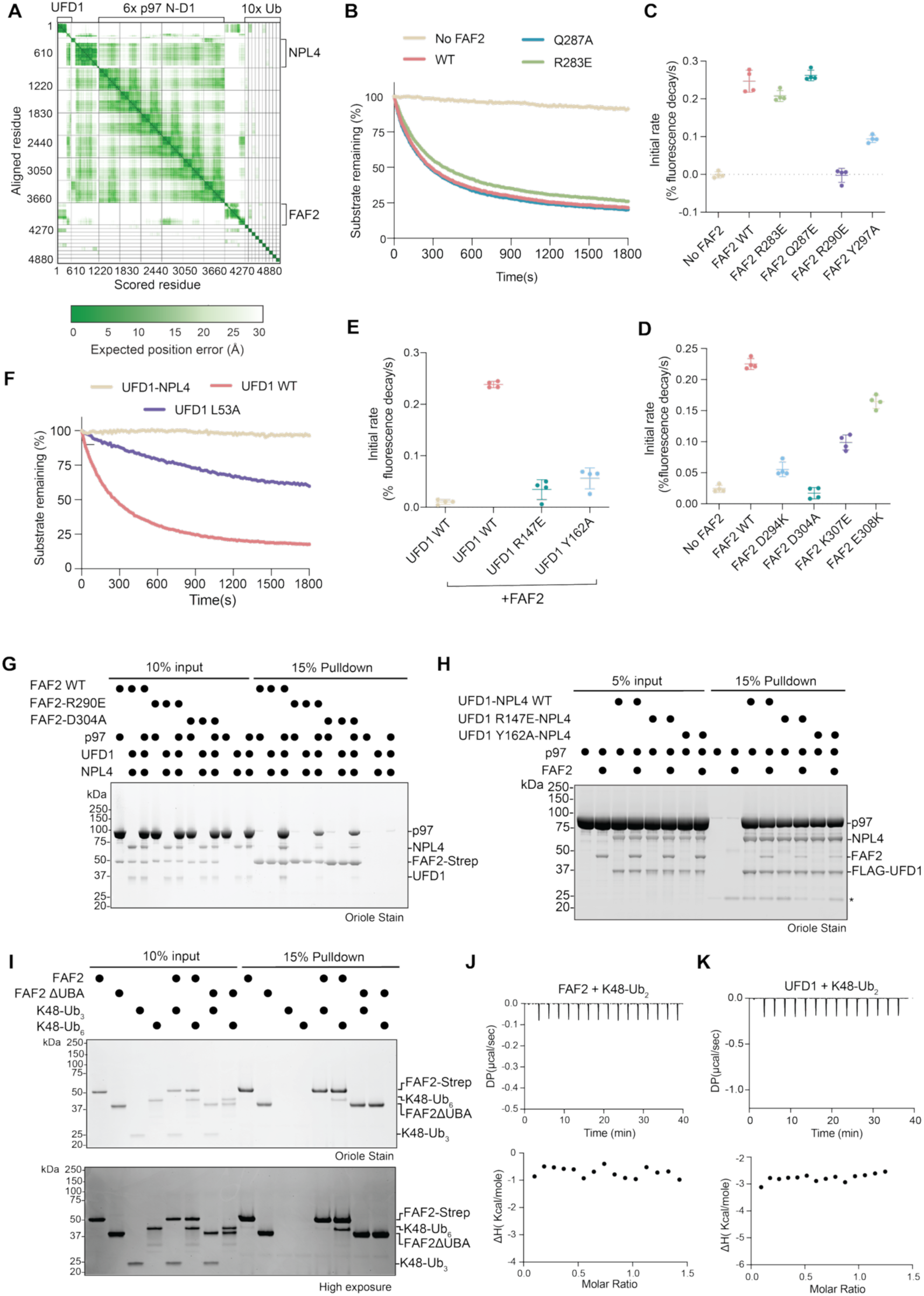
Analysis of FAF2, UFD1 and Ubiquitin interactions. A) PAE plot for the AlphaFold3 model of p97(ND1)-UFD1-NPL4-FAF2 with 10 ubiquitin molecules. (B) Unfolding activity of FAF2 mutants predicted to abrogate interaction with proximal Ub, measured in presence of UFD1-NPL4. (C-D) Comparison of substrate unfolding kinetics for FAF2 mutants predicted to impair UFD1 and ubiquitin interactions. (E) Comparison of substrate unfolding kinetics for UFD1 mutants predicted to impair FAF2 interactions. (F) Effect of disrupting the UFD1–Ile44 proximal ubiquitin interface on substrate unfolding activity measured in presence of FAF2. (G) Streptavidin pulldown assays assessing interactions between FAF2 (WT and indicated mutants) and the p97–UFD1–NPL4 complex. (H) FLAG pulldown assays assessing binding of wild-type or mutant Flag-tagged UFD1–NPL4 complexes to p97 and FAF2. (I) Pulldown assays testing interactions between strep-tagged FAF2 and K48-linked ubiquitin chains (Ub_3_ and Ub_6_) in the absence of the UBA domain. Full-length FAF2 was included as a control. (J) Isothermal titration calorimetry (ITC) analysis of FAF2 binding to K48-linked diUb. Representative thermograms (top) and binding isotherms (bottom) are shown. (K) Isothermal titration calorimetry (ITC) analysis of UFD1 binding to K48-linked diUb. Representative thermograms (top) and binding isotherms (bottom) are shown. In B and F, traces were normalized to fluorescence intensity at time 0 and substrate-only controls. For (B-F), data is shown as mean ± SD for n = 4 technical replicates.

**Figure EV5:**
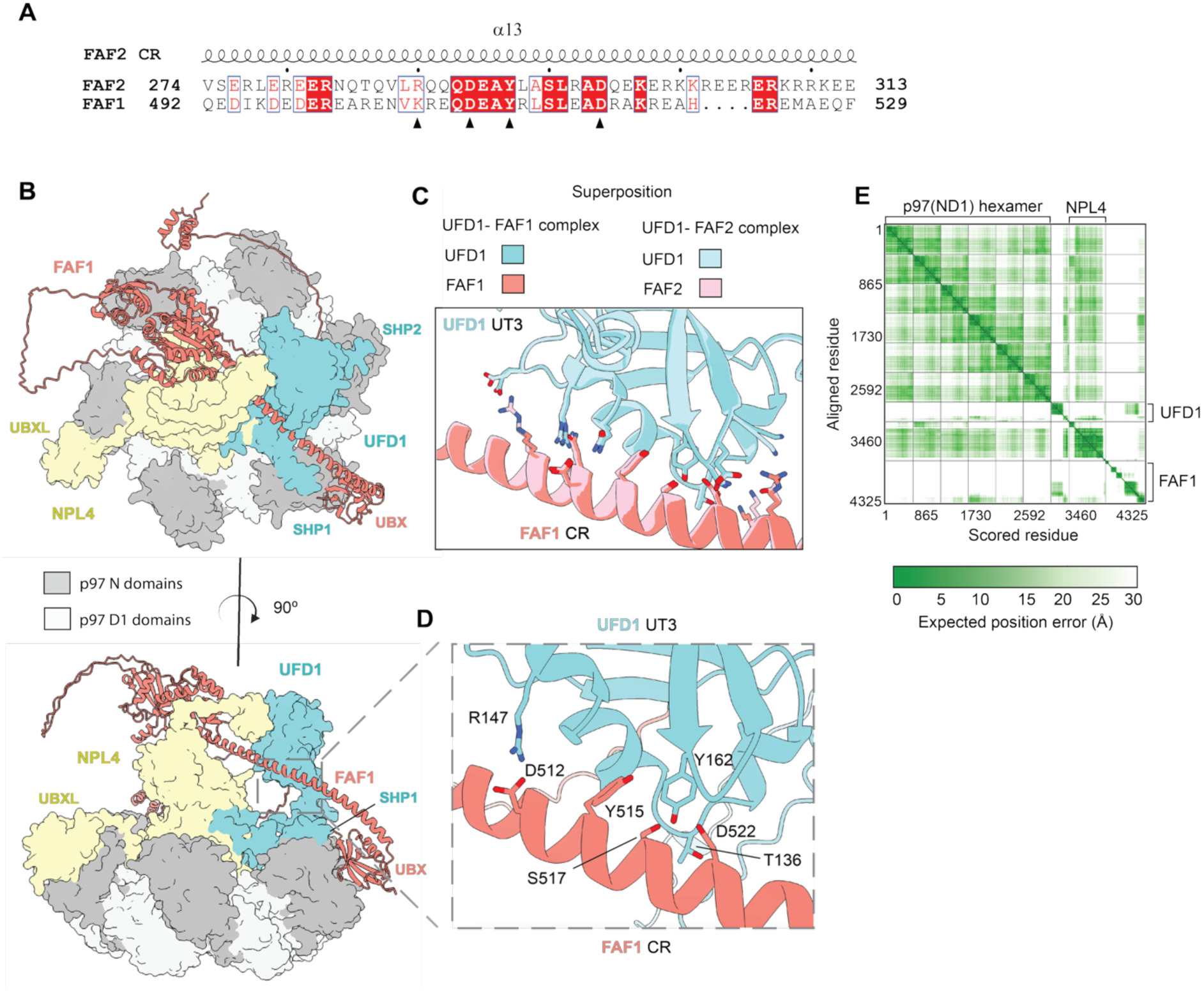
FAF1 engages p97–UFD1–NPL4 via a binding mode similar to FAF2. (A) Sequence alignment of the CR of human FAF1 and FAF2. Black triangles denote residues previously shown to contribute to UFD1 binding and enhanced unfolding activity. (B) AlphaFold3-based structural model of the p97(ND1) hexamer bound to the UFD1– NPL4 complex and FAF1. p97 N domains are shown in dark grey and D1 domains in light grey, NPL4 in yellow, UFD1 in teal, and FAF1 in salmon. The top panel shows a top view of the hexamer, highlighting the position of FAF1 relative to the NPL4 UBX domain, UFD1 SHP domains, and p97 N domains. The bottom panel shows a side view of the hexamer, illustrating how FAF1 binds the p97 N domain while bridging the UFD1–NPL4 complex. (C) Structural superposition of FAF1–UFD1-NPL4-p97 and FAF2–UFD1-NPL4-p97 AlphaFold models, revealing a conserved binding mode. Key interface residues conserved between FAF1 and FAF2 are shown as sticks. (D) Enlarged view of the FAF1–UFD1 interface showing specific residue–residue contacts. Key interacting residues are labelled. (E) PAE plot for the AlphaFold3 model of p97(ND1)-UFD1-NPL4 with FAF1.

**Figure EV6:**
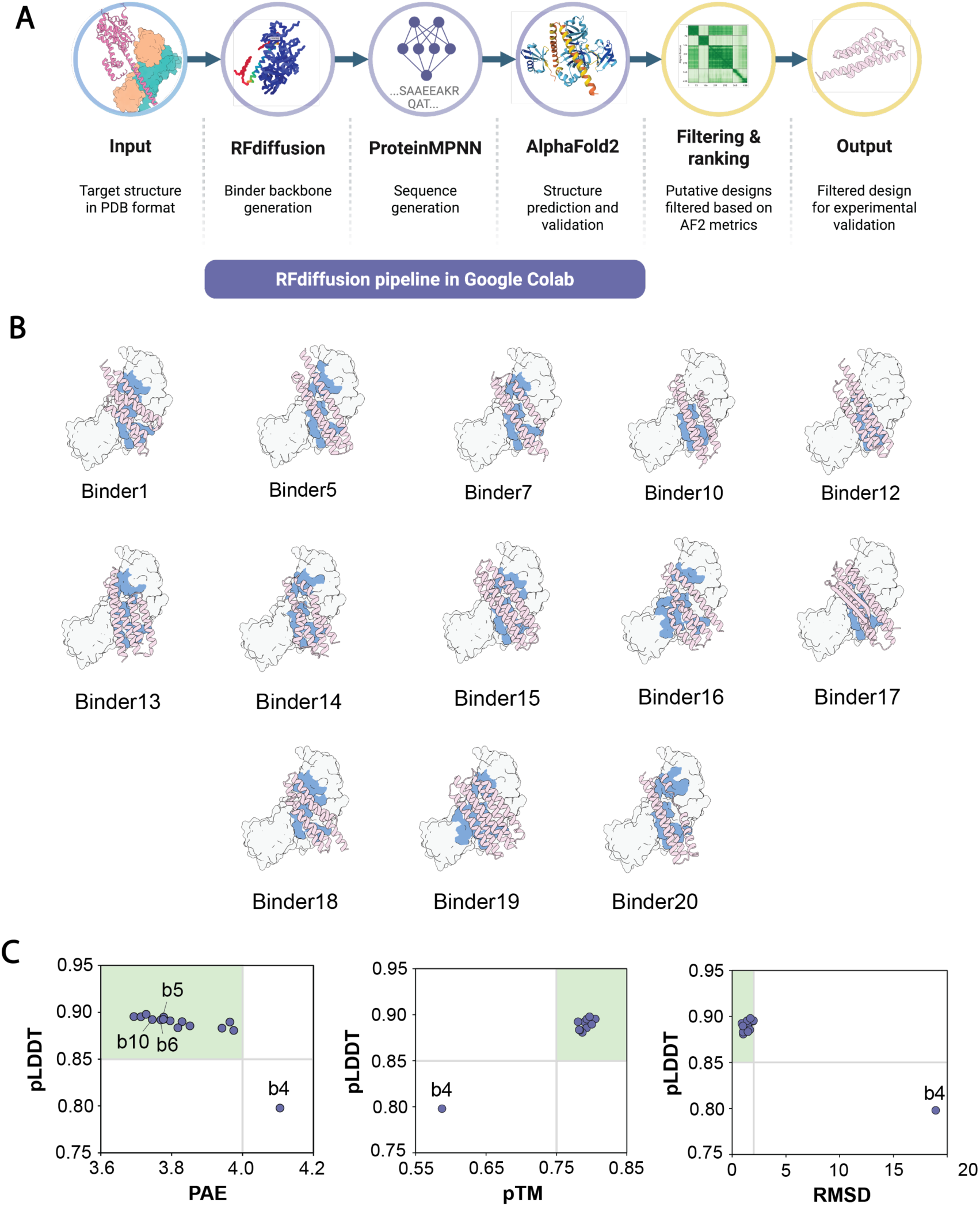
An overview of UFD1 binder design pipelines. (A) Schematic representation of the RFdiffusion pipeline used to design p97 activators. Given an AlphaFold prediction of the p97-UFD1-NPL4-FAF2-diUb complex as a target structure, RFdiffusion generates binder backbones conditioned upon the FAF2 helix. These backbone structures are then run through Protein MPNN to design amino acid sequences encoding the predicted structures, followed by model validation with initial guess AlphaFold2. The resulting designs are then filtered based upon AF2 output metrics. (B) Designs of potential p97 activators (pink) binding to UFD1 and a K48-linked ubiquitin dimer (shown in grey surface model). The residues on UFD1 and ubiquitin predicted to make the strongest and most stable contacts with the binder are shown in blue. (C) AF2 metrics of potential activators chosen by experimental validation are shown here. Regions corresponding to accepted designs are shown in green (PAE <4, pLDDT >0.85, pTM >0.75, RMSD <2). Binder4 initially failed on all metrics (as shown), but subsequent remodelling of the sequence in ColabFold (Mirdita et al. 2022) using the RFdiffusion input model as a custom template to guide the prediction improved the metrics considerably (pLDDT = 0.86, pTM = 0.76).

**Figure EV7:**
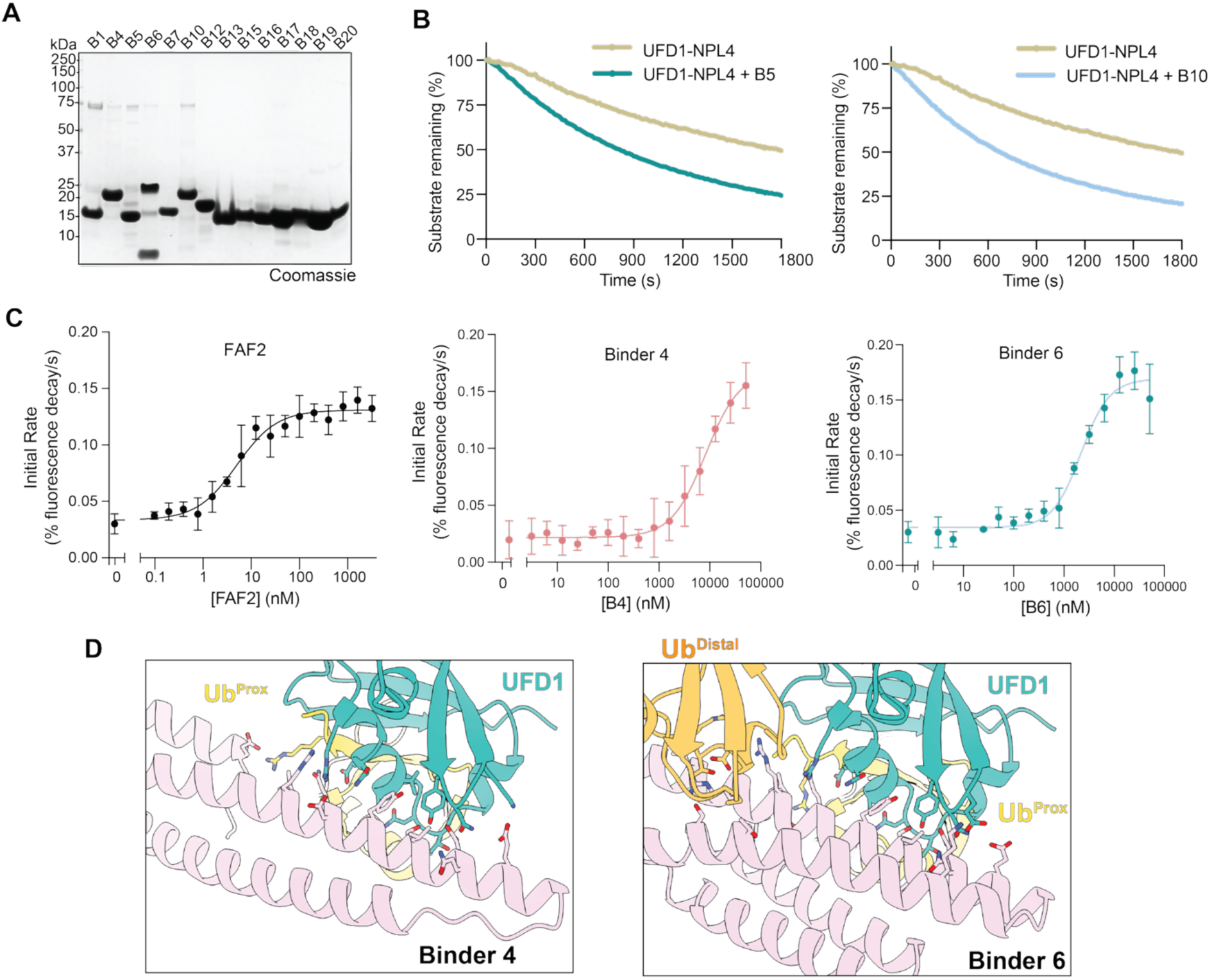
Characterization of *de novo* binders. (A) Coomassie-stained SDS–PAGE shows 14 purified *de novo* binders used for experimental validation. (B) Unfolding activity of p97-UFD1-NPL4 complex in presence of binder 5 (left) and binder 10 (right) shows modest activation of Ub^L^ substrate unfolding. Traces were normalized to fluorescence intensity at time 0 and substrate-only controls. (C) Initial rate of unfolding Ub^L^-Eos as a function of FAF2, binder 4 and binder 6 concentration. Initial rate was calculated from linear regression fitting of first ∼2 min of the unfolding trace. Data shown as mean ± SD for n = 3 technical replicates for FAF2, n=4 technical replicates for binders. (D) Structural models of UFD1 (teal) and two ubiquitin molecules bound to Binder 4 and Binder 6. Ubiquitin molecules are shown in yellow (proximal) and orange (distal), with interface residues highlighted as sticks.

## Notes

### Competing Interest Statement

The authors have declared no competing interest.

